# Reliable prediction of short linear motifs in the human proteome

**DOI:** 10.64898/2026.03.04.709504

**Authors:** Rita Pancsa, Erzsébet Fichó, Zsofia E. Kalman, Csongor Gerdán, István Reményi, András Zeke, Gábor E. Tusnády, Laszlo Dobson

## Abstract

Short linear motifs (SLiMs) are small interaction modules within intrinsically disordered regions of proteins that interact with specific domains, and thereby regulate diverse biological processes. The limited information in SLiM definitions leads to frequent false positive hits in computational and experimental SLiM identification methods. Here, we refined the annotations of known motif classes to create a high-quality training dataset, and developed SLiMMine, a deep learning-based method to identify SLiMs in the human proteome. Relying on protein embeddings, SLiMMine outperforms similar approaches in identifying functional SLiMs, while eliminating ∼80% of the pattern matching-based hits as false-positives. Furthermore, it can uncover novel SLiMs that do not fit known motif classes purely based on sequence features. Finally, narrowing the broad interactor-domain definitions of SLiM classes to specific partners enables more precise linking of predicted SLiMs to known protein-protein interactions. SLiMMine is available as a user-friendly, multi-purpose web server at https://slimmine.pbrg.hu/.

## Introduction

Short linear motifs (SLiMs) are protein segments typically composed of 3–10 amino acids that can interact with specific protein domains^1^. Their structural flexibility and limited number of specificity-determining residues allow them to *de novo* emerge in proteins through evolution^2^. SLiMs usually mediate weak, dynamic and transient interactions, often through coupled folding and binding, which enables them to function in cooperative regulatory systems. SLiMs are the characteristic interaction modules of intrinsically disordered regions (IDRs), which provide approximately half of the solvent-accessible surface of the human proteome^2,3^, therefore, millions of SLiMs are expected to regulate the diverse biological processes of human cells^4^. They are implicated in various diseases, ranging from disease-associated mutations that abolish the motif function^5^ to pathogenic effector proteins that mimic SLiMs to hijack host-cell regulation^6^.

Despite the growing knowledge on SLiMs, their identification remains challenging. Both computational and experimental methods struggle due to the limited sequence information within these short segments, which complicates robust screenings and leads to high false-positive rates^7^. Because multiple steps are needed to confirm a motif’s functionality, the number of classified examples remains limited to only a few thousand^8^. In accord, current resources for SLiMs are also limited. The Eukaryotic Linear Motif (ELM) resource is richly annotated, and its curation process incorporates detailed structural, functional and evolutionary analyses^8^, but other SLiM sources (such as LMPID^9^ or UniProt annotations^10^) do not cover all these aspects.

Numerous attempts have been made to enrich the limited information stored in these databases, including high-throughput experimental and computational approaches. The proteomic peptide-phage display (ProP-PD) protocol identifies short linear motifs at the proteome scale. Initially developed for PDZ domains^11^, it has now been extended to other SLiM-binding bait domains, enabling high-throughput discovery of domain–SLiM interactions across the human proteome^12^. Complementing these experimental strategies, quantitative fragmentomics studies such as those by Gogl *et al.*^13^ have systematically measured tens of thousands of PDZ-binding motif (PBM)-PDZ domain affinities within the human interactome, providing a detailed landscape of PDZ binding specificity and promiscuity. In parallel, computational approaches can identify protein binding segments in disordered regions based on sequence profiles^14^, biophysical and energetic properties^15,16^ or protein embeddings^17^, find conserved segments^18^, apply contextual filters^19^, detect motifs based on AlphaMissense pathogenicity scores^20^ or prioritize those motifs of a protein, whose partner domains are present in the experimentally verified interaction partners of the protein^21^.

Here, we propose an accurate deep learning-based method for identifying SLiMs in the human proteome that largely eliminates the problem of abundant false-positive hits. We relied heavily on the manually curated ELM resource and manually refined and extended its annotations to construct a high-quality training dataset that represents a high added value. Using protein embeddings and various neural network architectures, we developed a predictive model that reliably identifies novel SLiM candidates. The accuracy of the method is demonstrated on diverse independent test sets and case studies. Furthermore, based on the new, more detailed annotations and available protein-protein interaction (PPI) data, we pinpoint the likely interaction partners of the predicted motif instances, and thereby also propose a likely interaction mechanism for a large number of human PPIs, as well as predict novel PPIs.

## Results

### Refinement of ELM class annotations to gain an optimal training set for developing a SLiM predictor

To train a SLiM predictor and to identify the most likely binding partners of the predicted SLiMs among known PPIs, we needed reliable SLiM data annotated along specific criteria. As a first step, we mapped mammalian motif instances to their human orthologs and adjusted the motif boundaries of human instances to comply with current UniProt^10^ sequences.

To obtain a training dataset that meets our pre-defined criteria, we have refined and extended the annotations for the ∼320 ELM motif classes relevant to human proteins. Relying on ELM class text descriptions, the biological context of the motif classes and related literature, we determined for each class if the motif-containing protein counterparts 1) are extracellular or intracellular, 2) if intracellular, what compartment they are expected to localize to, and 3) what their expected Gene Ontology^22^ (GO) biological processes and molecular functions are. In addition, whenever possible, we precisely defined the possible binding partners, i.e. the subset of human proteins carrying the specific binding domain that actually binds to the given SLiM class. This was required because ELM domain definitions are often not specific enough. For instance, the docking motif specific for JNK kinases (ELM class: DOC_MAPK_JIP1_4) was assigned to the generic kinase domain, which can be found in hundreds of human kinases. In our refined dataset, only the three human JNK kinases are listed as possible partners for this motif class. In other cases, broad definitions of the interacting domains could be changed to more specific definitions from InterPro^23^. Precise definition of the possible partners for each ELM class enabled us to determine the most likely binding partners of the predicted motif instances in later steps: for a predicted motif instance in any given protein, the overlap between the possible partners of the respective ELM class and the known binding partners of the protein (in the human interactome) provides the most likely binding partners. From now on, we refer to this modified version of ELM data as ELM_refined (Figure 1/A).

**Figure 1.**
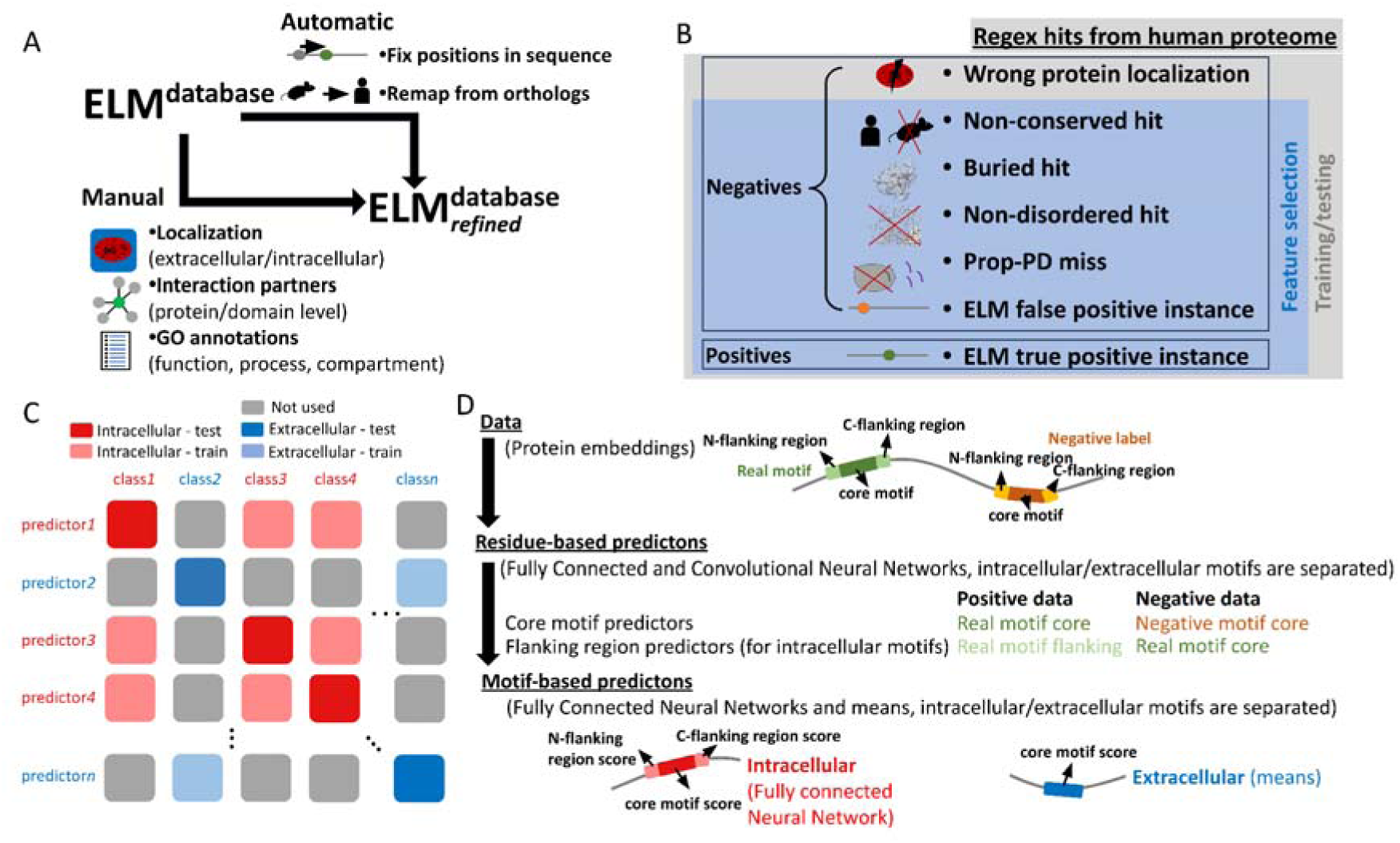
The training data, architecture and accuracy of SLiMMine. A) Automatic remapping of motif instances and manual refinement of ELM class annotations. B) Definition of labels used to train (and test) SLiMMine. C) Definition of predictors, training set and testing sets considering the motif class and the broad-level localization definition of the motif class. D) Prediction of SLiMs: Schematic view of how different motif regions are defined for the predictors (including residue-level and the final motif-level predictors).

### Assembling a reliable benchmark set for the training of SLiMMine

Classified ELM instances with the adjusted motif boundaries were used as positive data. Constructing a negative dataset is more challenging, as we cannot genuinely rule out regions without negative evidence, except for motifs labeled as false positives in ELM. As the latter dataset is very small, we defined a negative dataset for each motif class. To this end, we defined a set of criteria that could indicate if a given protein or region is unlikely to carry a true functional motif belonging to a given class. We scanned the human proteome for regular expressions (regex) defined in ELM and flagged all motif hits using the following criteria (these definitions may overlap). I) The protein localization is not appropriate for the motif class. II) The motif hit is not conserved in mammals. III) The motif hit is buried in a monomeric PDB^24^ structure. IV) The motif hit was not predicted to be disordered using AIUPred^25^. V) The motif hit, although tested, did not bind to the annotated partner domain in ProP-PD^12^ experiments. VI) False positive annotation in ELM. Each motif class has a different number of annotated instances (with experimental evidence) and motif hits in the proteome (based on how permissive the regex is). Using the flagged (likely false-positive) motif hits, five times more negative data were selected for each class than the associated positive data (Figure 1/B).

### Accurate prediction of Short Linear Motifs

Compared to the estimated number of SLiMs involved in cell regulation^4^, the number of experimentally verified motifs is very low, complicating the training and objective benchmarking of prediction methods. To overcome this problem, we created a separate prediction for each motif class. The training data includes all (positive and negative) motif instances from the same cellular milieu (intracellular or extracellular) that do not belong to the selected class, and then the predictor was benchmarked on instances of the particular class (Figure 1/C). This design ensures independence at the motif-class level: instances of the tested motif class are excluded from training. However, because individual proteins can contain motifs belonging to multiple classes, proteins may occur in both the training and test sets. We used a Gaussian mixture model for data augmentation, thereby increasing the number of positive instances to match the number of negative examples.

For all predictions, the input specifies the start and end positions of the candidate motif within a protein. For intracellular motifs, we first built residue-level predictors for core motifs (using the positive and negative motifs as input) and flanking regions (where the positive data consisted of the flanking regions of verified motifs, and the negative data consisted of the core motif regions) (Figure 1/D). In each case, a fully connected and convolutional Neural Network (NN) was used: while the core motif predictors have 87% accuracy (Figure 1/D), the flanking region predictors exhibit somewhat limited usability (62-63% accuracy). The lower accuracy is not unexpected, as motifs often stack on each other (therefore, the flanking and core regions may overlap), and different motif classes rely on flanking regions to a different extent for specific binding. By incorporating all residue-level prediction results into a fully connected NN, the prediction accuracy reached 94% for intracellular motifs (Figure 1/D). For extracellular motifs, we used only core motif predictions (Figure 1/C), and the final score is the mean of the residue-level prediction results, with 95% accuracy (Figure 1/D).

We conducted additional tests to objectively evaluate the performance of SLiMMine. Using the benchmark set, we assessed the performance of a classical approach that incorporates protein disorder (from AIUPred) and conservation (for more details, see methods). We also benchmarked AIUpred-binding^15^, a successor of Anchor^26^. Although it was not specifically developed for motif analysis, it is widely used, available as both a web server and a standalone program, and its authors have demonstrated its ability to identify functional motifs. We also assessed the performance of other methods predicting disordered binding segments, such as MoRFchibi 2.0^16^ and DeepDisoBind^14^. SLiMMine outperforms all other approaches on our benchmark set (Figure 2/A, SFigure 1). Notably, the classical conservation-disordered approach also performs well, however preparing the alignments for this task is a rather time consuming and laborious procedure. We also benchmarked SLiMSearch4^19^, allowing regular expression scanning and applying contextual filters. The server did not allow to search all motifs (for example if the motif regular expression yields too many hits) and it uses a different version of the human proteome, therefore we could not completely benchmark SLiMSearch4. On the remaining motifs and proteins listed by the server, SLiMMine filters (after applying 0.5 probability threshold) excluded more actual motifs, however at the same time eliminated negative cases much more successfully (SFigure 2). While the training/benchmark sets may be selected arbitrarily, to demonstrate the power of our method, we tested SLiMMine on further independent datasets: phosphomodulated motifs^27^, cyclin-docking motifs^28^, 14-3-3 binding motifs^29^, essential motifs required for normal cell proliferation^30^ and LMPID^9^ motif entries (Figure 2/B), and it confidently identified the vast majority of the associated motifs (73.6% of the 1177 tested motifs received a score of 0.5 or higher, and 40% of them 0.9 or higher). Notably, SLiMMine scores are not exceptionally high for Prop-PD hits^12^, but when filtered to retain only conserved motifs, the resulting distribution becomes comparable to previously defined positive sets (SNote 1, SFigure 3). In contrast, segments obtained by pattern matching of ELM regular expressions on monomeric structures, coiled coils (decoy sets) and disordered linker and terminal regions (neutral set) received low scores (Figure 2/B). The distributions of scores for monomeric structures and coiled coils show that SLiMMine does not falsely identify motifs within these folded regions, even though coiled-coils are disordered in their monomeric form and are therefore often cross-predicted with disordered regions^31^. The distribution of SLiMMine scores for disordered linker and terminal regions is uniform, consistent with regex hits being either true motifs or falsely identified ones in these regions.

**Figure 2.**
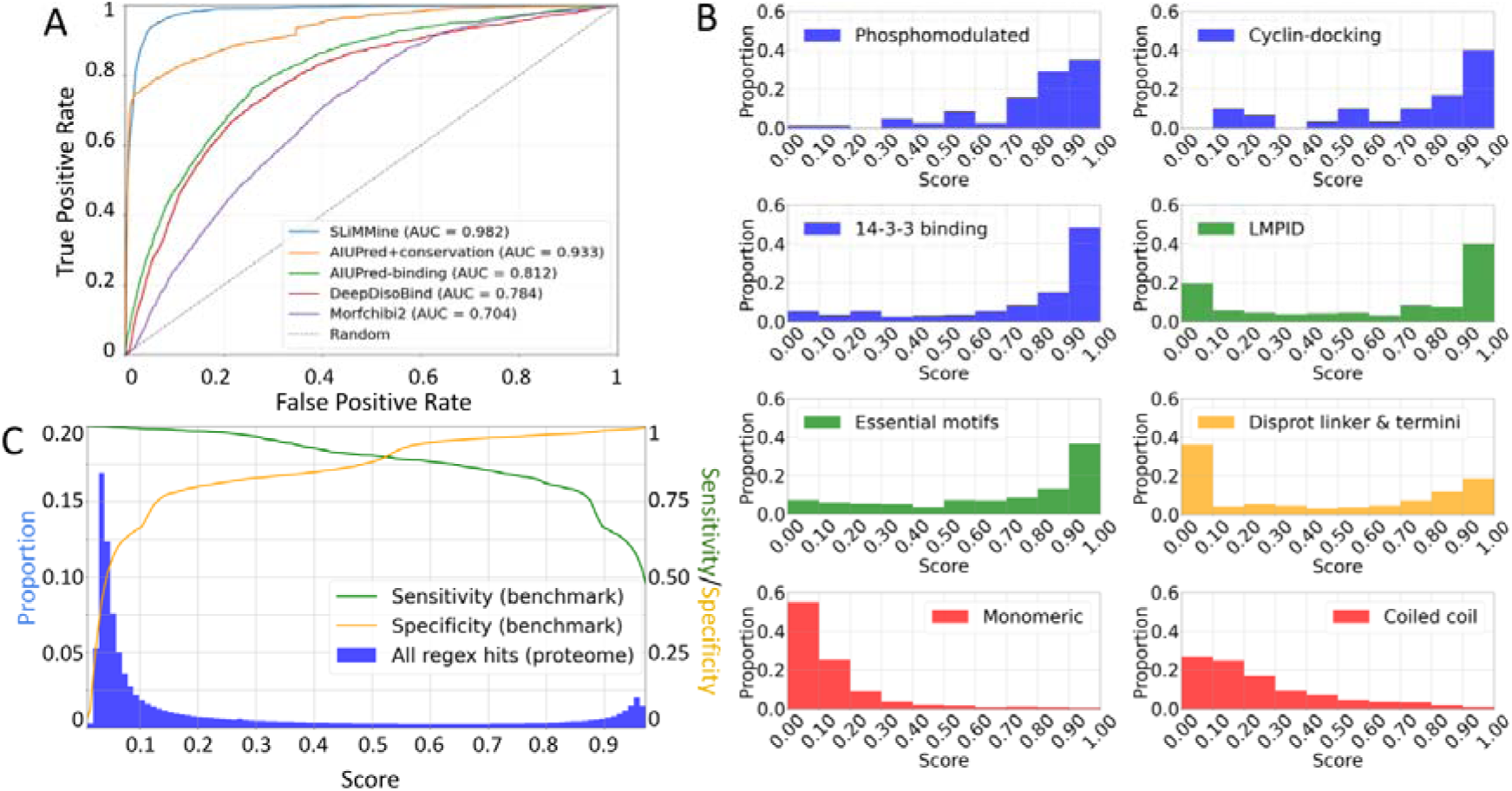
SLiMMine accurately predicts SLiM-mediated interactions across diverse datasets and in the human proteome. A) Receiver Operating Characteristics of SLiMMine, AIUPred-binding, DeepDisoBind, MoRFchibi 2.0 and a traditional prediction using conservation and protein disorder (AIUPred). B) Prediction score distribution on specific motif sets (blue: phosphomodulated motifs (n=83), cyclin docking motifs (n=30) and 14-3-3 binding motifs (n=263)), general motif sets (green: LMPID motif entries (n=276) and essential motifs (n=525)), a neutral set of disordered regions (yellow: ELM class regex hits within disordered linkers and termini (n=2032)) and decoy sets (red: ELM class regex hits within monomeric structures (n=1374) and coiled-coils (n=1092)). These sets are independent from training data, since motifs overlapping with ELM instances have been removed. C) Distribution of prediction scores for all motif regex matches in the human proteome (bars). Sensitivity and specificity values calculated on the benchmark set at varying score thresholds are indicated by lines.

**Figure 3:**
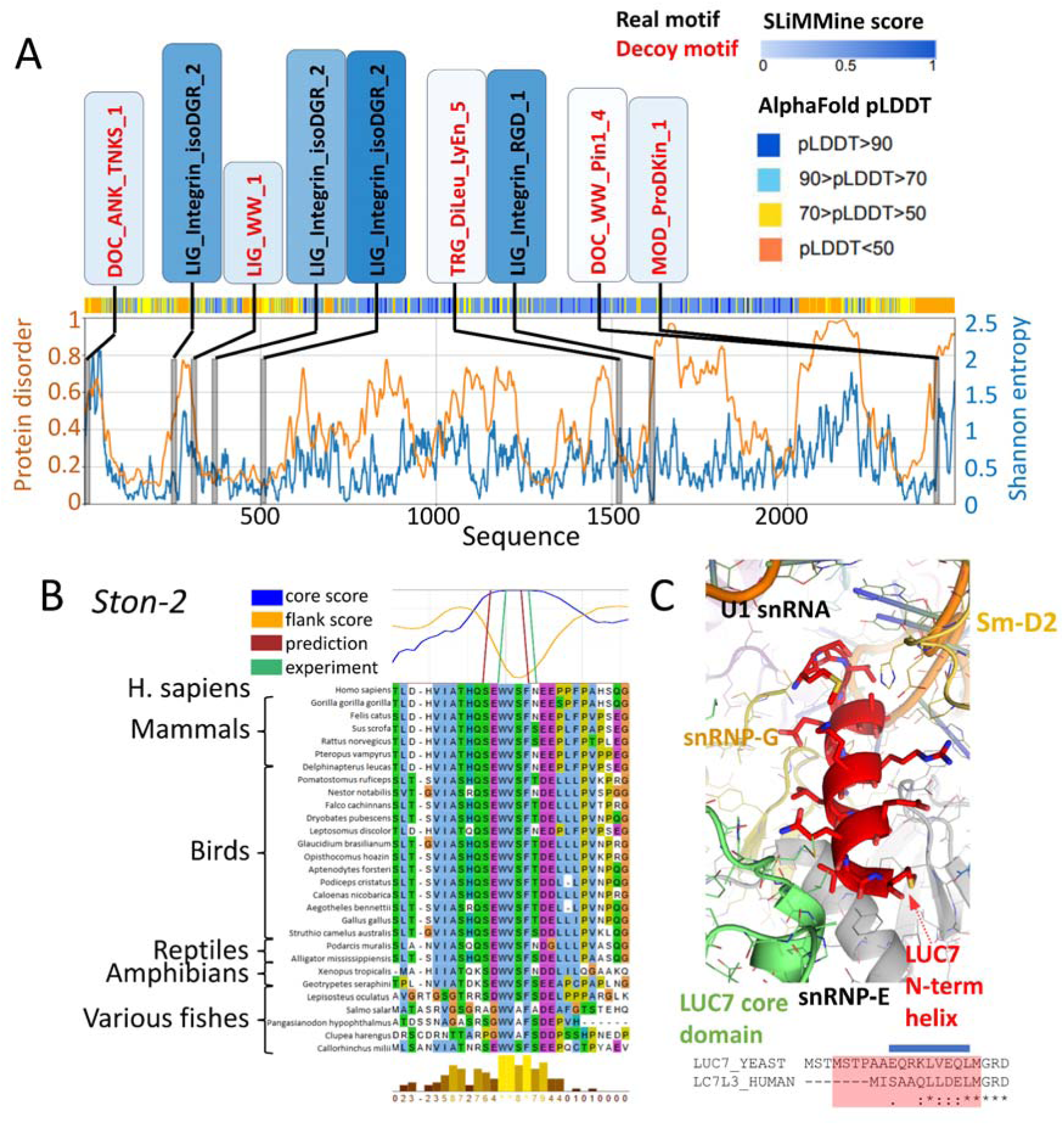
SLiMMine successfully identifies novel instances of known motif classes as well as yet undefined motifs. A) SLiMMine prediction results of known motif classes on Fibronectin using real and decoy motifs. AlphaFold pLDDT values, AIUPred output and residue conservation are also displayed. B) Class-independent motif prediction, alignment and protein conservation for a SLiM in the Stonin-2 protein, which binds to the alpha-ear domain of AP-2 complex subunit alpha-1. C) Structural snapshot of the N-terminus of yeast Protein LUC7 in the spliceosomal E complex (pdb:6N7R, in red), contacting multiple spliceosomal proteins (colored variously) as well as the U1 spliceosomal RNA (orange). The alignment with the N-terminus of the human Luc7-like protein 3 protein is shown below, with the red box highlighting the ordered amino acids observed in the crystal structure. The conserved segment, predicted as a class-independent motif by our algorithm, is indicated by a blue bar.

### Prediction of Short Linear Motifs across the human proteome

Considering all regex hits in the proteome, SLiMMine scores show a bimodal distribution (Figure 2/C, the reference distribution on the benchmark set is visible in SFigure 4). While there are hundreds of thousands of motif regex hits in a grey zone where specificity is lower, the proportion of hits is higher above the 0.9 threshold, where the specificity is 0.98. Altogether, we predicted 304,238 SLiM instances with a score of 0.9 or higher, and 696,329 SLiM instances with a score of 0.5 or higher, among known motif classes. This methodology could filter out 2,718,778 regex hits as likely false-positives (score < 0.5).

**Figure 4:**
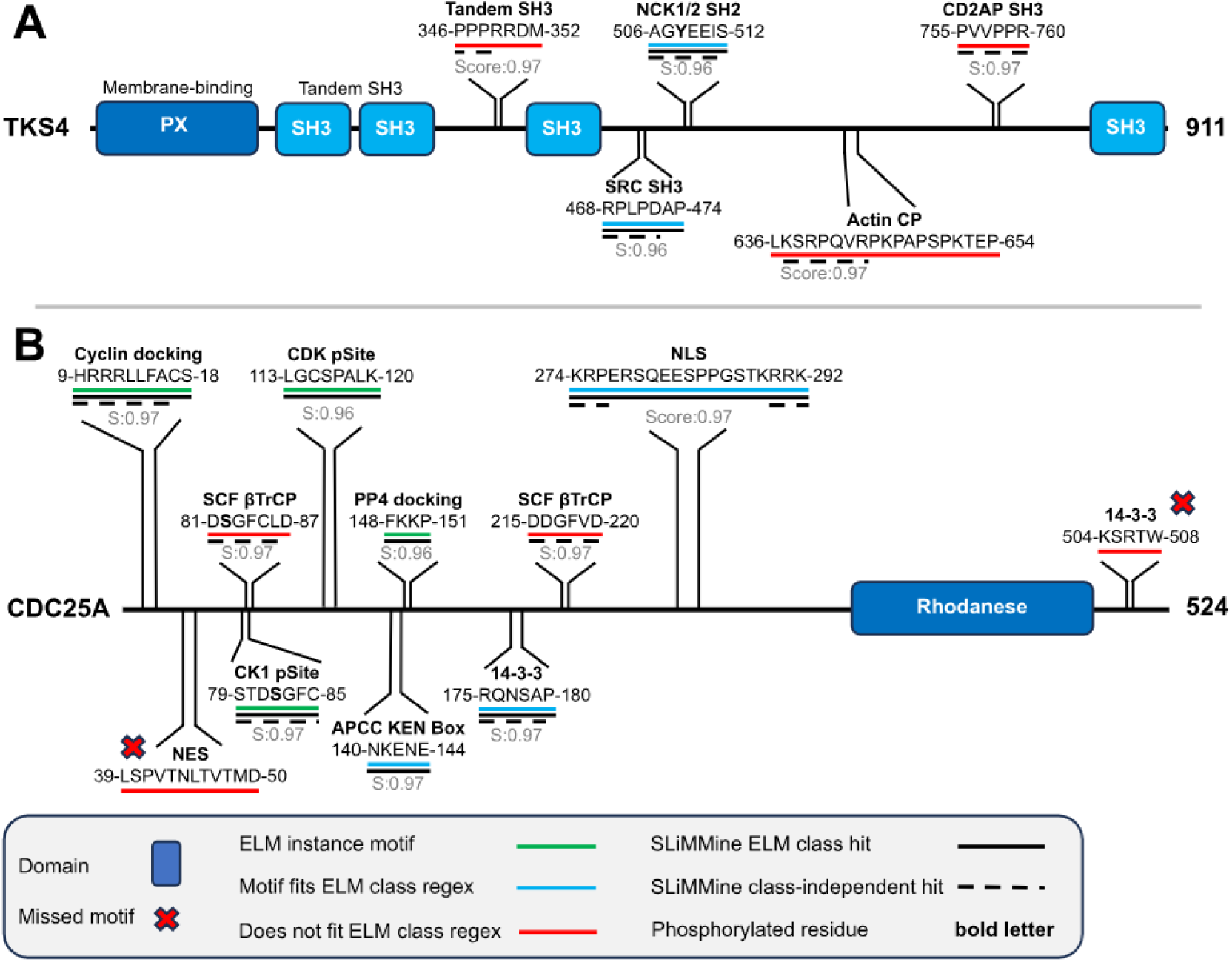
SLiMMine successfully identifies most known motifs of the human Tks4, CD2AP and Cdc25A proteins. A) The domains (blue boxes) and SLiMs (sequence bits) are indicated on the domain maps of Tks4 and B) Cdc25A. For each known motif, the respective binding domains/motif names, sequences, residue boundaries, ELM status (underscoring in different colors) and SLiMMine predictions (underscoring with black simple (ELM class predictions) and/or dashed (class-independent predictions) line) are indicated along with the respective SLiMMine Score (S).

The power of SLiMMine prediction is demonstrated on Fibronectin (*FN1*; UniProt: P02751), which contains multiple integrin-binding motifs. These are often located in loops of ordered domains, thus they are not necessarily disordered. In addition, the residue-level conservation value (using Shannon entropy) fluctuates along the Fibronectin sequence, making motif identification harder. Still, SLiMMine successfully identifies all experimental motifs, whereas decoy motif regex hits (meaning regex hits for intracellular motifs in extracellular or buried regions) were assigned with lower prediction scores (Figure 3/A).

To also showcase an intracellular example, in the Smaug1 RNA-binding protein (*SAMD4A*; UniProt: Q9UPU9), SLiMMine correctly identifies three of the four experimentally validated 14-3-3 binding motifs^32^, the nuclear export signal^33^ (NES) and the C-terminal PDZ-binding motif (Dean group, unpublished results) with very high prediction scores (>0.9), although none of these motifs were present in the training set (SFigure 5).

**Figure 5:**
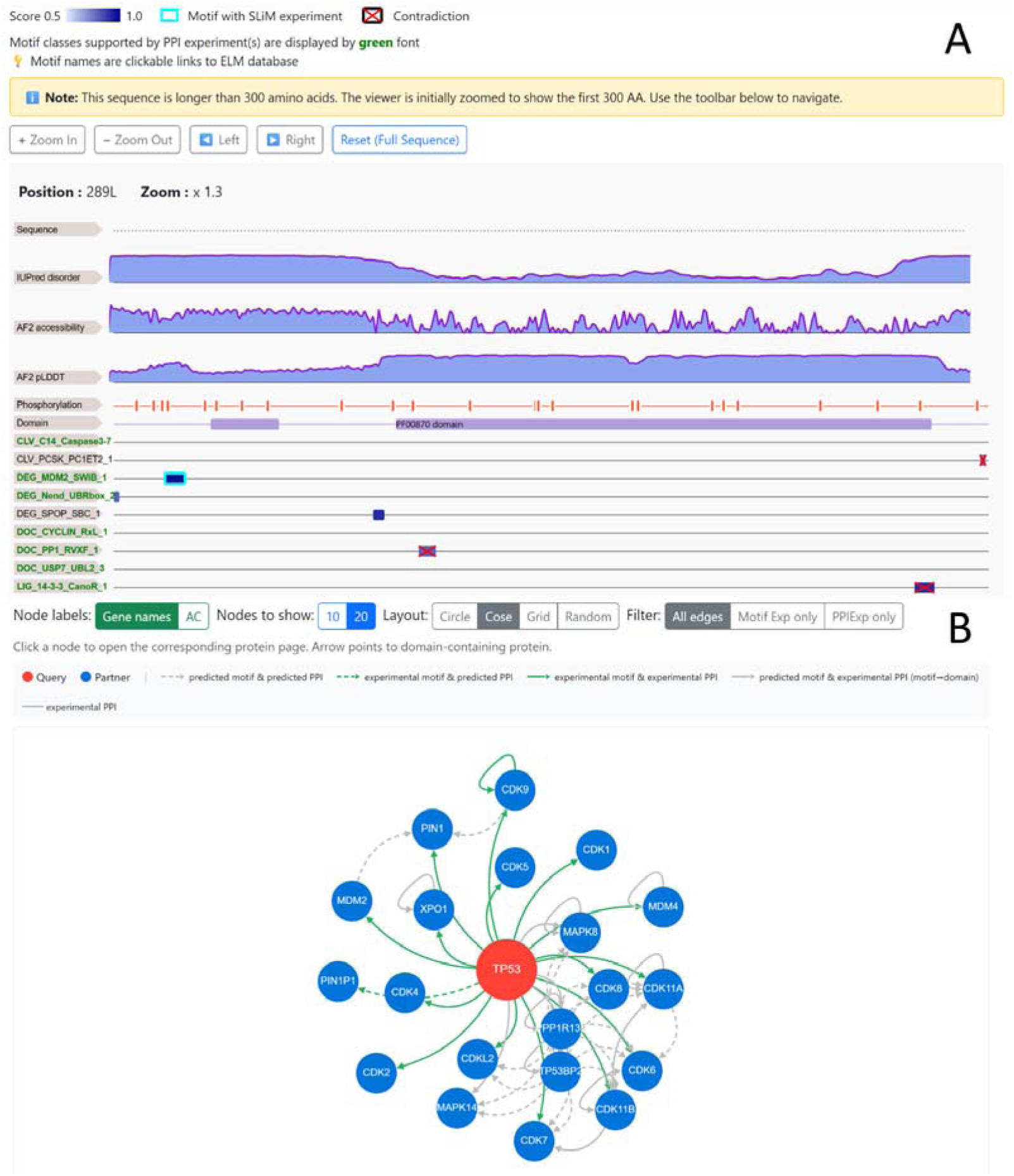
The SLiMMine web server represents an ideal tool for motif discovery. A) *Layout of the sequence viewer and B) the Protein interaction viewer.*

These results underscore that SLiMMine can reliably identify functional instances of the already defined motif classes while efficiently reducing the associated false-positive hits.

### SLiMMine enables the discovery of novel motifs that may not adhere to existing class definitions

SLiMMine also enables the discovery of novel SLiMs class-independently based on sequence context. The reason behind this is that SLiMMine operates on protein sequence embeddings rather than raw sequence patterns, so it does not memorize specific known motifs. Instead, it learns general properties characteristic of SLiMs as encoded in the embedding space. Notably, as the flanking region predictors could not be reliably calibrated for extracellular motifs, this functionality is only available for intracellular motifs.

While the core motif predictor in SLiMMine identifies longer regions as favourable for motifs, flanking region prediction opens prospects for defining novel motifs without specifying their start and end positions. We rationalized that if the core motif scores are high and the flanking region scores are low in the very same region, and in addition, the flanking region scores increase before and after this segment, the analysed region is probably a SLiM. Considering the constructed benchmark set, SLiMMine predicted class-independent motifs (i.e. when the start and end positions were not given) with 0.98 specificity (and a lower 0.56 sensitivity) on motifs of 3-9 residues in length, with a standard error of 1.91 residues on motif boundaries (considering the start and end positions as defined by the regex). Applying SLiMMine with these thresholds on the full human proteome across all possible lengths, identified 32,501 short protein regions. 1,713 of these motifs do not overlap with any sequence matches to ELM-defined regular expressions, and the ones that do overlap may also be novel motifs that just overlap with the given (degenerate) regex match by chance. These therefore represent particularly interesting candidates for motifs that are not captured by currently defined ELM motif patterns. Notably, class-independent motifs precisely map to over a thousand experimentally validated motifs from ELM, and several motifs from PDB and LMPID that were not part of the training set (SFigure 6). For example, in Stonin-2 (*STON2*; UniProt: Q8WXE9), SLiMMine correctly identified the experimentally verified alpha-ear domain-binding motif^34^ stored in LMPID, even though it was not part of the training set (Figure 3/B).

A powerful feature of our class-independent detection is that it can also identify atypical motif variants not covered by standard regular expressions. As an example, the N-terminus of Dynein 2 intermediate chain 2 (*DYNC2I2*; UniProt: Q96EX3) can engage no less than 3 dynein light chain molecules in the dynein tail, forming 3 successive beta-sheets, according to the structure pdb:6RLB^35^. Here, our method successfully identified the 3rd segment as a novel motif, although its sequence “HVDAQVQT” does not respect the canonical [KR]xTQT pattern. SLiMMine also finds the conserved, unconventional 14-3-3 binding motif in CREB-regulated transcription coactivator 1 (*CRTC1*; UniProt: Q6UUV9), evidenced by the structure pdb:7D8H^36^, despite its complete lack of arginine/lysine residues (QYYGGSLP).

The flexible N-terminal region of the human Luc7-like protein 3 (*LUC7L3*; UniProt: O95232) was predicted to contain a novel motif that is conserved in its yeast homolog Protein LUC7. In the crystal structure of the *S. cerevisiae* spliceosomal E complex (pdb:6N7R^37^), this segment forms a short helix, binding simultaneously to U1 spliceosomal RNA, as well as proteins snRNP-E, snRNP-G and Sm-D2 (Figure 3/C).

In all, SLiMMine can successfully, and, regarding sequence boundaries, very precisely identify SLiMs that were not present in its training set and may not even adhere to the SLiM definitions it was trained on.

### Case studies

The accuracy of SLiMMine is also demonstrated on some full-length proteins that have multiple validated SLiM-mediated interactions covering a wide variety of motif classes. In the multi-domain podosome scaffold protein, Tks4 (*SH3PXD2B*; UniProt: A1X283), SLiMMine correctly identified the experimentally validated tandem SH3-binding motif^38^, actin capping protein (CP)-interacting motif^39^ and CD2-associated protein (CD2AP)-binding motif^40^ as class-independent motifs, although none of them are present in ELM. While ELM stores 9 class variants for SH3-binding motifs, the tandem SH3-binding motif is not among them^41^. The CD2AP-binding motif of Tks4 does not completely fit the strict regex of the corresponding ELM class (class: LIG_SH3_CIN85_PxpxPR_1, regex: Px[PA]xPR) because it has a valine in the middle^40^. However, results of peptide array screens support that the three SH3 domains of CD2AP have slightly different recognition specificities, and some of them also accept valine at this position^42^. The identified CP-interacting motif also does not fit the ELM class definition (LIG_ActinCP_CPI_1) due to a slight difference, however, subunits of the actin CP heterodimer were consistently identified as Tks4 binding partners by immunoprecipitation assays in five cancer cell lines^39^, and therefore the proposed motif is most likely functional. On top of the interesting class-independent hits, SLiMMine also pinpoints a Nck1/2 SH2-binding motif at phosphosite Y508, enabling the interaction between Tks4 and Nck1/2^43^. The SH2 domain of Src recognizes the same phosphorylated Tyr (Y508), while its SH3 domain binds to a Pro-rich peptide within Tks4^44^ that SLiMMine also identified as a high-scoring hit. Although our institute has a long history in studying Tks4, we do not know of any other approved motifs in it^38–41,44^. Therefore, although not encountered during training, SLiMMine confidently identified all five known motifs of Tks4 (Figure 4/A). Of the ten class-independent motif hits within the 911 residues long Tks4 (which cover <10% of the sequence) five match already validated motifs, while the remaining five could be yet uncovered motifs. The interaction partner of Tks4, CD2AP is also presented as a case study (SMaterial, SFigure 7).

The M-phase inducer phosphatase 1 Cdc25A (*CDC25A*; UniProt: P30304) is often used to illustrate how the versatility of SLiMs can regulate the life cycle and functioning of a protein^1,4^. It has 11 well-documented motifs, of which four are annotated ELM instances^8^ (the cyclin-docking motif, a CK1 and a CDK phosphorylation site and the PP4 phosphatase docking motif). All four are successfully identified by SLiMMine (although in the case of the cyclin-docking motif and CK1 phosphorylation site, the motif boundaries are slightly shifted compared to the ELM instance (see SNote 2 and SFigure 8 for explanation)). Three other motifs that fit the respective ELM class regex definitions are also all identified (the KEN-box degron^45^, a canonical 14-3-3-binding motif^46^ and the nuclear localisation signal^47^). The two copies of the SCF-beta-TrCP E3 ligase-specific degron^48^ do not exactly fit the respective ELM regex, still, they are both found as class-independent hits. Only two Cdc25A motifs were missed: the NES^47^ that does not fit ELM NES class definitions and is too long to be identified as a class-independent hit, and a second, non-canonical 14-3-3 binding site centered around T507^46^ that is in a non-disordered region (AIUPred score < 0.4). In all, SLiMMine could identify most known motifs of Cdc25A as high-scoring hits (Figure 4/B).

Additional case studies, including E3 ubiquitin-protein ligase RNF13, Smaug1, CD2AP, LanC-like protein 2, and Early E1A from Human adenovirus 5 further illustrate the applicability of SLiMMine across diverse biological contexts (SNote 3, SFigure 5, SFigure 7, SFigure 9, SFigure 10).

### SLiMMine predictions can be linked to protein-protein interaction (PPI) data and explain underlying binding mechanisms

After predicting novel instances for the ELM-derived SLiM classes in the human proteome, we also created a network of potential interactors using the newly defined sets of partner proteins created for the classes (refined based on the ELM database’s descriptions and associated literature as described in the the 1st section of the Results). For each predicted motif, experimentally verified PPIs of the given protein (from PDB (through 3did^49^), IntAct^50^ and BIOGRID^51^) were screened to identify partners that were among the predefined partners of the respective motif class. This procedure yielded a list of likely SLiM-mediated interaction partners for each protein with predicted motifs and, at the same time, suggested underlying interaction mechanisms and interacting regions for more than 50 000 human PPIs (SNote 4l, SFigure 11), for which no such data were previously available.

### SLiMMine comes with a user-friendly web server facilitating motif discovery

SLiMMine is shared with the research community through a modern, easy-to-use web resource. To cover all possible use cases, the server provides three different options. Users can search for I) any protein in the human reference proteome, II) predefined ELM classes or III) custom regular expressions. The first option leads to the respective protein entry page with all predicted motifs displayed, while the latter two options provide a list of proteins that contain sequence matches to the selected ELM class/custom regex, along with the associated SLiMMine prediction scores.

All search results can be further inspected on the entry page, which has five main blocks: General, motif (viewer), motif details, protein interactions (viewer) and protein interaction details. The General section displays general information about the protein, including subcellular localisation, domains, and statistics.

The motif viewer (Figure 5/A) was designed to summarize all relevant information about the predicted motifs along the sequence and to display evidence-based filters that help the user decide whether the motifs could be functional. At the top of the panel, measures related to protein disorder are shown. Each motif class is shown below in a separate line. The different colors and frames of the boxes representing the motifs correspond to different evidence supporting the actual motif. For example, motifs with contradicting evidence (e.g., a phosphodependent motif without experimental phosphorylation evidence^52^) are crossed. Notably, when we defined these contextual filters, we found that they are unlikely to rule out functional motifs (SFigure 12) and they often help to pinpoint functional motifs among multiple hits (SFigure 13). Below the viewer, there is a table detailing supporting (or contradicting) evidence on the motifs.

The protein interaction network (Figure 5/B) is largely based on the partner lists for the SLiM classes and displays the best annotated SLiM-mediated interactions for the entry protein. Predicted and experimentally supported motifs, as well as protein interactions can be viewed and filtered. The table below details all possible SLiM-mediated interactions of the entry protein (all predicted SLiM classes with all their possible partners listed in the refined SLiM class annotations) with supporting evidence also displayed.

## Discussion

SLiMMine is an alignment-free approach for predicting SLiMs. It is optimized for scenarios in which motif boundaries are known, for example, after scanning sequences with regular expressions derived from motif databases. The model evaluates whether the given region exhibits properties consistent with functional SLiMs. Although the method was trained and optimized on human protein data, we expect it to perform similarly well on other mammalian proteins.

A major obstacle during the development was the limited availability of experimentally validated motif data. Positive samples could be retrieved from ELM^8^. Defining true negatives was more challenging, as the absence of a motif annotation does not imply the absence of function (see SNote 5). However, emerging high-throughput experimental approaches are expected to eliminate this limitation in the long term^12,53,54^. As coverage of such experiments increases - especially for viruses and other underrepresented taxa - these data will provide valuable benchmarks for evaluating and retraining models such as SLiMMine. The current version of SLiMMine has not been tested for SLiMs in various pathogenic species, yet initial benchmarking shows promising results (see SNote 6, SFigure 10 and SFigure 14). Notably, SLiMMine was trained and tested on conserved motifs, so the prediction of fast-evolving viral mimicry motifs^55^ may need further considerations^8^.

SLiMMine classifies more than 80% of the proteome-scale sequence matches of ELM-derived regular expressions as false positives. However, it is important to note that even among the predicted positives, erroneous hits may still be present. One contributing factor to false predictions is that some motif classes have permissive regular expressions with higher pattern probabilities and therefore occur tens of thousands of times within the human proteome. In such cases, sequence-based prediction alone is insufficient to distinguish functional instances from coincidental matches. To mitigate this issue, we introduced refined and extended ELM annotations that incorporate contextual constraints, such as phosphorylation dependence, subcellular localization, or restriction to transmembrane proteins. For example, motifs that are known to function only when phosphorylated are explicitly annotated as such. The SLiMMine web resource highlights these contextual annotations. Notably, the human proteome is not uniformly annotated, therefore, it is up to the users to decide whether they accept the neural network score or rely on the provided supporting or contradicting evidence.

Beyond identifying candidate SLiMs, SLiMMine provides additional biological insight by proposing likely interaction partners for predicted motifs, and thereby enables functional hypothesis generation. The combined information can serve as a powerful input for both computational and experimental studies, and can substantially reduce the search space in large-scale interaction modeling efforts, such as AlphaFold-based domain–motif screening approaches^56^, or guide the design of targeted validation assays, such as peptide pull-downs, mutational analyses, or cell-based interaction assays.

An important extension of SLiMMine is the class-independent motif prediction module, which seeks to identify SLiM-like regions without predefined boundaries. This idea is motivated by the observation that many SLiMs exhibit “island-like” conservation patterns^57^: a short core region shows strong functional constraint, while flanking residues are more variable but still contribute to binding specificity or regulation. By evaluating the relationship between the predicted motif core and its surrounding residues, SLiMMine can highlight regions without defining the start and end position of a segment. Notably, if multiple motifs overlap, the interpretation of flanking regions becomes ambiguous. In all, the class-independent approach can be valuable for identifying novel motif classes and atypical binding modes that are currently underrepresented in curated databases.

SLiMMine can serve as a starting point for addressing biologically relevant questions. Many regulatory proteins contain multiple SLiMs that engage several binding partners, leading to cooperative or competitive interaction networks^58^. Multivalency can increase effective binding affinity and enable dynamic assembly of signaling complexes^1^. In addition, clustering of multiple SLiMs may contribute to liquid–liquid phase separation, in which networks of weak, multivalent interactions drive the formation of dynamic biomolecular condensates that spatially organise signaling and regulatory processes^59,60^. SLiMMine is well-suited to exploring this complexity, as it can identify multiple candidate motifs within the same protein and link them to distinct or shared binding domains.

SLiM prediction has important implications for understanding human diseases. Although disease-causing mutations are less frequent in disordered regions, many disrupt SLiMs^20,61^. Loss (or gain) of a single motif can rewire interaction networks, leading to aberrant signaling, mislocalization, altered protein turnover or stability^62^. By highlighting predicted motifs and their potential interaction partners, SLiMMine can aid interpreting missense variants that fall outside classical functional domains.

In all, with its accurate predictions, SLiMMine represents a large step forward in identifying and understanding short linear motifs in human proteins. We plan to update SLiMMine when new data become available (i.e. when new motif class definitions can be extracted from novel, annotated SLiM databases or derived from large-scale experimental SLiM discovery datasets) and extend it to proteins of other organisms. In addition to predicting novel instances of already defined SLiM classes, SLiMMine could also identify ∼32,500 class-independent SLiMs in the human proteome. Among these, we identified several experimentally validated motifs that were not part of the training data and may not fit existing motif definitions, providing strong support for the applicability of our method. Furthermore, reliable motif predictions from SLiMMine and the refined partner annotations for hitherto known motif classes enabled rendering likely binding mechanisms and interacting regions to a large number of human PPIs, as well as highlighting novel motif-mediated PPIs with high probability. The associated user-friendly, multi-purpose web server is a powerful tool with the potential for taking motif discovery endeavours to the next level.

## Methods

### Manual curation of data from the ELM Resource

We have refined and extended the annotations for the 319 motif classes in the ELM resource that are relevant to human proteins. For each class, we determined if the motif-containing protein counterparts 1) are extracellular or intracellular, 2) if intracellular, what compartment they are expected to localize to, and 3) what their expected GO^18^ biological processes and molecular functions are. Although ELM has its own GO annotations, those are mixed GO terms, meaning either relevant to the motif- or domain-containing counterparts of the interactions (or both), so they would have been problematic to use. We also identified the potential binding partners for the given ELM class - that is, the subset of proteins containing the relevant binding domain that has been shown to interact with that particular SLiM type. For this, either the same (usually PFAM) domain identifier was used as in ELM (if further specification of the partners was not required), or a more specific domain identifier from InterPro^23^, or the list of UniProt ACs of the possible interaction partners within the human canonical proteome. We also defined if a motif is only present in proteins with a particular domain or in transmembrane proteins. Last but not least, we catalogued all motif classes that 1) require phosphorylation for binding or 2) result in phosphorylation (as a product of binding), so that available proteome-scale phosphorylation data could be used to support or contradict the functionality of the associated predicted instances (SData 1, SNote 7).

### Automatic processing of ELM data

Motif instances and classes were downloaded from the Eukaryotic Linear Motif Resource in November 2024^8^. We checked if instance boundaries in *H. sapiens, M. musculus,* and *R. norvegicus* proteins were altered by mapping them to the respective canonical sequences in UniProt^10^ (release: 2025 January). If the motif could not be found at the given position, but the sequence still carried the indicated peptide in a single copy within a disordered region (AIUPred^25^ > 0.4), we modified the region boundaries accordingly. In addition, we aligned *M. musculus* and *R. norvegicus* proteins with their *H. sapiens* orthologs (sharing the same gene name) and transferred motif boundaries from the orthologous sequences onto the human counterparts (SData 2).

### Defining motif labels

The positive set was defined using all true positive human instances in ELM, extended with those mapped from orthologous species. The negative set contains false positive human instances from ELM. As this set is minimal, we scanned the human proteome for the regular expressions stored in ELM, and defined the following labels. I) Residues of the putative motif are also part of a PDB monomeric structure (defined by SIFTS^63^), and their mean relative solvent accessible area is lower than 0.09. II) The regular expression defining the putative motif had lower than 0.3 conservation within the multiple sequence alignment (MSA) in a set of Chordata species (SData 3) (see Conservation). III) If the localization of the protein rules out the functionality of the putative motif (considering cytoplasmic, nuclear and extracellular motifs). IV) The putative motif is in an ordered region (average AIUPred score across the putative motif residues was below 0.3). V) The respective SLiM-binding domain was used as bait in ProP-PD experiments, but the peptides carrying the putative motif were not found as positive hits^12^.

The ELM Resource categorizes motifs into classes (based on interacting with the same binding site of a specific partner domain and the function), and each class contains multiple experimentally validated motif instances. For each class, the positive set included all positive instances from that class. The negative set, on the other hand, included five times more regex hits associated with the same class than the positive data. The negative set was selected to ensure a roughly equal representation of the following cases: I) ordered putative motifs, II) non-conserved putative motifs, and III) putative motifs from proteins with an inappropriate subcellular localization. In addition, less populated negative cases, including false-positive ELM instances, motifs buried within monomeric structures and ProP-PD^12^ negative results (where the peptide carried the motif pattern but it was not bound by the corresponding SLiM-binding domain), were included (SData 4). Notably, this ratio may vary depending on the regular expression definition: if a motif pattern only occurred in a limited number of proteins, fewer negative samples were used.

### Conservation

We defined motif conservation in two ways. If a regular expression was available, we used a set of Chordata species proteomes (SData 3) and searched orthologs for human proteins using BLAST^64^ (the sequence with the highest sequence identity (above 20%) was selected for each species, if it covered at least 60% of the human sequence), and then aligned the selected sequences with ClustalΩ^65^. We searched for the regular expression in the orthologous proteins. Since aligning disordered regions is challenging, we allowed 50 residues deviations in the alignment for regular expression hits. Non-conserved motifs were defined with <0.3 conservation, while conserved motifs were defined with >0.7 conservation across orthologs. To visualize entropy for the Fibronectin example, we simply calculated the Shannon entropy at each position in the alignment.

### Train and test sets

For each motif class, we defined a broad-level localization definition (extracellular/intracellular, SData 1). We created a separate predictor for each class (SData 5), where the test set included all instances from a given class. In contrast, the training set contained instances from all the other classes with the same broad-level localization. This way, it was possible to perform independent tests for each ‘class’ predictor. Of the 319 manually refined ELM classes, 290 contained at least one suitable positive instance for benchmarking and were therefore used to construct predictors; the remaining 29 classes and the respective reasons for their exclusion are detailed in SNote 8.

### Architectures and training

The goal of the method is to take a short protein segment (i.e. start and end position in a sequence, that is usually defined by a regular expression) and predict whether that segment exhibits SLiM-like characteristics, that is reflected by a score between 0 and 1. We created multiple predictors for each class using different regions of the motif and different neural network (NN) architectures. First, we separated extracellular and intracellular motifs. In both cases, we defined a core motif predictor (prediction on the actual motif, relying on the previously defined dataset). In addition, for intracellular motifs, we developed a flanking region predictor (for extracellular flanking regions the NN did not converge), defined as five neighbouring residues on both sides of the actual motif. For the flanking predictor, the negative data consisted of the actual motif (see Figure 1/D). The training data is based on the prottrans_t5_xl_u50 embeddings^66^ of residues in the motif, and these predictions are performed on the individual residues. For each predictor, we randomly selected 10% of the positive and negative data and used recursive feature elimination to select the top 100 features. Notably, for this step, we omitted all negative instances for which the negative annotation was derived from the incorrect localization of the protein, as it seemed that, in such cases, the embedding-based predictors learnt the localization of the motif and did not consider any other biophysical or biological features. Nevertheless, feature selection was performed 50 times, and features appearing in 50% of the cases were selected for all prediction types (i.e. intracellular core, intracellular flanking and extracellular). As the training set contained five times more negative data, we augmented positive data by adding Gaussian noise to it (on the level of embedding vectors) with a 5% standard deviation. Then, we used both fully connected and convolutional NNs to predict core and flanking regions. In the case of intracellular motifs, a fully connected NN was used to predict the final result for a given segment, using the residue-level core and flanking NN predictions as input. For extracellular motifs, the final prediction is the mean of the outputs of the residue-level fully connected and convolutional NNs. The architectures of these predictors are visible on SFigure 15. The results of the residue-level predictions on the training and test sets are available in SData 6-13 (SData 6: ‘core’ inside motifs, convolutional NN, SData 7: ‘core’ inside motifs, fully connected NN, SData 8: flanking inside motifs, convolutional NN, SData 9: flanking inside motifs, fully connected NN, SData 10: ‘core’ outside motifs, convolutional NN, SData 11: ‘core’ outside motifs, fully connected NN, SData 12: Statistics on motif classes, SData 13: Overall statistics). Final prediction results of the motifs on the training and test sets are available in SData 14-17 (SData 14: inside motifs, SData 15: outside motifs, SData16: Statistics on motif classes. SData 17: Overall statistics). To verify that our method is not overfitting, we used the same procedure with mixed labels too. As expected, the NNs could not converge and produced ∼45-55% prediction accuracy.

### Independent test sets

The constructed train/test sets were also used to benchmark traditional conservation/disorder-based motif prediction. We used AIUPred^25^ results (calculating the mean over the residues within the motif) and conservation score from Chordata species (see “Defining motif labels”), then the mean of these two values defined the score of the traditional approach. We also compared SLiMMine with AIUPred-binding^15^, DeepDisoBind^14^, and MoRFchibi 2.0^16^(SData 18, Figure 1/A). We also compared SLiMMine performance with SLiMSearch 4 contextual filters (SFigure 2, SData 19).

We also used additional motif data to test the robustness of our method. Cyclin-docking peptides were collected from Örd M. *et al.*^28^, where ∼100,000 peptides were assayed for relative binding strength to cyclins from five human cyclin families (D, E, A, B, and F) using quantitative intracellular binding and large-scale tiled peptide screening. We relied on Supplementary Data 2 of the cited paper, from which only high-confidence hits were retained; when the authors identified a motif match, we refined the peptide boundaries to the detected motif. We collected phospho-modulated motif interactions^27^ identified via a phosphomimetic proteomic peptide–phage display (ProP-PD) assay using a library covering ∼13,500 phospho-serine/threonine sites in intrinsically disordered regions. We relied on Table S5 of the cited publication and accepted only high-confidence non-redundant peptide hits corresponding to the wild-type form. We used a compilation of 14-3-3-binding motifs from Egbert CM *et al.*^29^; however, we considered only a four-residue environment around the phosphorylated positions (listed in Table S2 of the cited paper). We also assessed the performance of SLiMMine on a list of essential human motifs required for normal cell proliferation based on large-scale mutation screening^30^ (obtained from the Source data of Figure 2 of the cited paper). We further tested SLiMMine on human LMPID^9^ motif entries supported by experimental evidence (SData 20).

We defined a decoy set by searching motif candidates with regular expressions in the ordered monomer test set also used for benchmarking Anchor^15^ and in coiled-coil regions predicted by four different methods (from Supplementary Table 3 of Kalman *et al.*^67^). In addition, we analyzed disordered linker and terminal regions from DisProt^68^ as a neutral set (covered by the functional annotation categories: “flexible C-terminal tail”, “flexible N-terminal tail”, “entropic chains”, and “flexible linker”) by pattern matching of ELM class regular expressions for motif candidates (SData 21). In the above described datasets, we only considered hits that did not overlap with any known ELM instance to ensure independence from the training/test sets. Prediction results on these sets are available in SData 20-21.

### Class-independent motif prediction

Using the constructed benchmark set, we evaluated multiple score cutoffs to define the most reliable class-independent motif prediction. Using intracellular motifs of length 3-9 residues from the training and test sets, we benchmarked the correspondence between the final, the residue-level core and the residue-level flanking scores. SData 22 contains the sensitivity, specificity, and extent to which the region boundaries of the observed and predicted motifs differ for these thresholds. We achieved the best overall result when the score from the flanking region predictors on the core motif divided by the score from the flanking region predictors on the flanking region is below 0.5, and the final motif prediction score is above 0.9 (SData 23).

### Prediction of SLiMs in the human proteome

We scanned the human proteome with the regular expressions of the 319 ELM motif classes that are relevant to human proteins. Based on motif localization (intracellular or extracellular), we applied the appropriate predictors to each match, using the start and end positions defined by the regular expression. Motif scores were calculated as the mean probability across all final scores generated by the selected predictors (SFigure 16, SData24).

For class independent prediction, we screened the human proteome for all segments of length 9, 7, 5, and 3. When a segment was identified as a motif, overlapping peptides were excluded from further analysis.

### Protein-protein interaction data

For all predicted motif instances, we considered the list of annotated, possible partners of the respective SLiM class as potential interactors (SData 1). If an interaction was validated in BIOGRID^51^, IntAct^50^ (with physical association or stronger annotation (MI:0915)), or it was included in the PDB (accessed via 3did^49^), ELM or LMPID, we accepted it as an experimentally verified protein-protein interaction. The resulting subnetworks are displayed on the SLiMMine web server entry pages.

### Web server for SLiMMine

SLiMMine is implemented as a web application using Laravel 12, PHP 8.4, and MySQL 8.0. The platform integrates multiple predictive models, combining machine learning outputs with AlphaFold2-derived pLDDT confidence scores^69^, AIUPred disorder probabilities^25^, and UniTmp^70^ transmembrane topology annotations. Query performance is optimized through database-level pagination and pre-indexed tables, enabling efficient access to large datasets.

The interactive visualization module uses the FeatureViewer JavaScript library with jQuery 3.6 and Bootstrap 5.3, ensuring responsive behaviour across devices. Batch data export is supported in tab-separated format, including the underlying prediction probabilities. Benchmarking across the complete human proteome (>20,000 entries) demonstrated fast response times, achieved through optimized SQL statements and server-side caching.

### Visualization

Visualization of alignment on Figure 3/B was prepared with JalView^71^, protein structure on Figure 3/C was visualized using ChimeraX^72^.

### Statistics and Reproducibility

All ELM data was used for training and benchmarking. During benchmarking, any data that overlapped with the training data were excluded from the evaluation. Multiple independent benchmark datasets were used to evaluate the method and to demonstrate consistent performance across diverse data types and experimental conditions. Test sets were held back from training. Randomization was only performed during feature selection, where random 10% of the training data was used 50 times.

Data analysis was performed using Python 3.10.12 (https://www.python.org/) and NumPy 1.24.0 (https://numpy.org/).

## Supporting information

SNote

## Data availability

The numerical source data underlying all graphs are provided with this paper as Supplementary Data files at https://figshare.com/articles/dataset/34330041^73^. Summary of tables is available in the Supplementary Material.

## Code availability

**Codes** for slimmine are available at: https://github.com/laszlodobson/SLiMMine and https://zenodo.org/records/21931868^74^.

## Acknowledgment

This work builds on two decades of effort behind the Eukaryotic Linear Motif (ELM) resource. We thank Prof. Toby Gibson for his leadership and all the ELM annotators for their invaluable contributions.

## Author contribution

L.D. designed the study; R.P., G.E.T and L.D. raised funds; R.P, Z.E.K., A.Z, and L.D collected the data and developed the method; R.P., A.Z. and L.D conducted the analyses, E.F., C.G., I.R. and G.E.T. built the web resource; R.P. and L.D. wrote the paper. All authors contributed to the drafts and gave final approval for publication.

## Funding

The project was implemented with the support from the National Research, Development and Innovation Fund of the Ministry of Culture and Innovation, financed under the K-146314, FK-142285 and PD-146564 funding schemes granted to G.E.T, R.P. and L.D, respectively. R.P is a holder of the János Bolyai Research Fellowship of the Hungarian Academy of Sciences (BO/00174/22).

## Declaration of interest

E.F. is an employee of Cytocast Hungary Kft. Furthermore, E.F. owns equities or stocks of the company. All other authors declare no competing interest.

## Notes

https://zenodo.org/records/18785791

https://zenodo.org/records/18786455

### Summary of Updates

Updated evaluation of the method. Amended manuscript.

