## Supplementary material for "Reliable prediction of short linear motifs in the human proteome": SNote

[1]Institute of Molecular Life Sciences, Research Centre for Natural Sciences, Magyar Tudósok Körútja 2, Budapest 1117, Hungary
[2]Department of Bioinformatics, Cytocast Hungary Kft, Petőfi Sándor utca 5/A, Budapest 1052, Hungary
[3]Faculty of Information Technology and Bionics, Pázmány Péter Catholic University, Práter utca 50/A, Budapest 1083, Hungary
[4]Department of Artificial Intelligence, Faculty of Informatics, ELTE Eötvös Loránd University, Pázmány Péter sétány 1/C, Budapest 1117, Hungary
[5]Department of Biology, University of Padua, Via Ugo Bassi, 58/B, 35131 Padua, Italy
[6]Department of Bioinformatics, Semmelweis University, Tűzoltó u. 7, Budapest 1094, Hungary
[†]These authors have contributed equally to the work.

### Table of content

List of Supplementary data files and figures 1
SNote 1: Limitations of high-throughput experimental datasets 3
SNote 2: Regular expression yielding overlapping motifs 3
SNote 3: Further cases of SLiMMine successfully detecting experimentally verified motifs 4
SNote 4: Protein-protein interaction data 4
SNote 5: Dataset construction 5
SNote 6: Pathogenic motifs 6
SNote 7: Manual annotation of SLiMs 6
SNote 8: Motif classes without any positive experimental instances 7
Supplementary Figures 9
References 20

**List of Supplementary tables, data files and figures**

Supplementary Data files (available at https://figshare.com/articles/dataset/34330041):
**SData 1:** Refined manual annotations of ELM resource motif classes (localization, binding partners and gene ontology)
**SData 2:** ELM motif instances subjected to automatic processing (transfer from ortholog, fixing region boundaries)
**SData 3:** List of species used for creating alignments and calculating conservation
**SData 4:** Train and test sets of SLiMMine
**SData 5:** Motif classes used to benchmark different predictors

**SData 6:** Prediction results on the core ‘inside’ motifs (convolutional NNs, residue level prediction)
**SData 7:** Prediction results on the core ‘inside’ motifs (fully connected NNs, residue level prediction)
**SData 8:** Prediction results on the flanking regions of ‘inside’ motifs (convolutional NNs, residue level prediction)
**SData 9:** Prediction results on the flanking regions of ‘inside’ motifs (fully connected NNs, residue level prediction)
**SData 10:** Prediction results on the core ‘outside’ motifs (convolutional NNs, residue level prediction)
**SData 11:** Prediction results on the core ‘outside’ motifs (fully connected NNs, residue level prediction)
**SData 12:** Summary statistics for residue level predictors (separated based on SLiM class used for testing)
**SData 13:** Summary statistics for residue level predictors
**SData 14:** Prediction results on ‘inside’ motifs (motif level prediction)
**SData 15:** Prediction results on ‘outside’ motifs (motif level prediction)
**SData 16:** Summary statistics for motif level predictors (separated based on SLiM class used for testing)
**SData 17:** Summary statistics for motif level predictors
**SData 18:** Comparison of motif prediction methods
**SData 19:** Comparison of SLiMSearch and SLiMMine
**SData 20:** Further independent test sets (cyclin-binding motifs, phosphodependent motifs, 14-3-3 binding motifs, essential motifs required for normal cell proliferation and LMPID motif entries)
**SData 21:** Further independent test sets (monomeric structures, coiled-coils and DisProt linkers/termini)
**SData 22:** Sensitivity, specificity and motif boundary statistics for class-independent predicted motifs at different thresholds
**SData 23:** SLiMMine class-independent motif prediction results on the benchmark set with selected thresholds
**SData 24**: Predicted motifs in the human proteome

Supplementary Figures:
**SFigure 1:** Comparison of precision-recall curve of SLiMMine and other methods.
**SFigure 2:** Comparison of SLiMSearch filters and SLiMMine predictions.
**SFigure 3:** SLiMMine performance on Prop-PD dataset.
**SFigure 4:** The performance parameters of SLiMMine on the benchmark set.
**SFigure 5:** SLiMMine finds almost all experimentally validated motifs of Smaug1.
**SFigure 6:** SLiMMine class-independent motif prediction finds multiple experimentally verified motifs.
**SFigure 7:** SLiMMine successfully identifies most known motifs of the human CD2AP.
**SFigure 8:** Overlapping and shifted motif detection may affect the visualization of ELM instances in SLiMMine.
**SFigure 9:** SLiMMine finds the nuclear localization signal in LanC-like protein 2 as a class-independent motif.
**SFigure 10:** SLiMMine successfully identifies the motifs employed by the viral Early E1A protein.

**SFigure 11:** SLiMMine predictions explain a non-negligible fraction of validated human PPIs.
**SFigure 12:** The effect of contextual contradictions on experimentally verified motifs.
**SFigure 13:** Context filters work in the Piezo2 receptor.
**SFigure 14:** SLiMMine performance on motifs of pathogens.
**SFigure 15:** The Neural Network architectures of SLiMMine.
**SFigure 16:** Use of predictors during benchmarking and proteome-wide prediction.

**SNote 1: Limitations of high-throughput experimental datasets**

We noticed that Prop-PD[1] results sometimes produce low scores. The most likely reason is that ELM[2] employs a rigorous annotation process and focuses on conserved motifs; however, not all motifs are uniformly conserved, if conserved at all. ProP-PD often tests interactions between domains and peptides that would never meet spatiotemporally in the cell; therefore, even if they bind in the experiment, the detected interaction might not be valid from the perspective of cell function. Also, some sticky peptides can produce nonspecific interactions that may not even involve the SLiM-binding pockets of the bait domains. Furthermore, the 16 residue long peptides from Prop-PD likely include extended flanking regions that may confound prediction. Therefore, we searched for the regular expression within the prey peptides known to bind the SLiM-binding domain used as bait. To further ensure that these peptides represent true functional hits, we filtered them for conservation: the identified regular expressions were subjected to conservation analysis (see Conservation section in the Methods). Although this procedure reduced the number of motifs in the analysis, SLiMMine scores are markedly higher on this filtered dataset (SFigure 3).

**SNote 2: Further cases of SLiMMine successfully detecting experimentally verified motifs**

The modular scaffold CD2-associated protein CD2AP (*CD2AP*; UniProt: Q9Y5K6) is the interaction partner of the Tks4. Interestingly, 3 of the 7 class-independent SLiMMine motifs overlap with three short Pro-rich peptide segments that can be bound by the Endophilin-A1 and Cortactin SH3 domains[3]. These SH3 domains do not have their specific motifs defined in ELM. However, the Endophilin-A1 SH3 domain prefers P+RPP motifs[4] (+ stands for positively charged residues), and the first peptide, which showed the strongest binding to this SH3[3], has a "PKKPP" segment that strongly resembles the preferred motif. Intriguingly, a SLiMMine-predicted class-independent motif exactly covers this segment (SFigure 7). Also, based on the sequence motifs of Cortactin SH3 partners collected by Roger J Daly[5], the optimal motif can be derived as +Px[VIAP]Px+P, where + stands for positively charged residues and x for any residue. Only the first and second Pro-rich peptides could bind to the Cortactin SH3[3], and both contain segments that strongly resemble this motif. SLiMMine also identified these as class-independent motif hits. Furthermore, SLiMMine could identify the actin CP-binding motif of CD2AP[6] (an ELM instance) and a proposed AP2-binding motif[7] that fits the regex of the respective LIG_AP2alpha_1 class (SFigure 7). In all, SLiMMine confidently identifies the known motifs of CD2AP, furthermore, four of the seven class-independent predicted motifs overlap with the validated motifs, and one with a predicted 14-3-3 binding motif (by 14-3-3-Pred[8]) centered on S458 that is a known phosphosite (CD2AP interacts with 14-3-3 proteins[9]).

LanC-like protein 2 is a peripheral membrane protein that associates with cellular membranes upon myristoylation and relocalizes to the nucleus when demyristoylated[10]. Truncation of the N-terminal region results in a homogeneous intracellular distribution.

Analysis using cNLS Mapper[11] identifies both a monopartite nuclear localization signal (NLS; Thr4–His12) and a bipartite NLS (Lys7–Gly37); however, neither conforms to the regular expressions stored in ELM. Notably, SLiMMine successfully detects two class-independent motifs within the N-terminal region (Met5–Lys10 and Glu20–Pro27), which overlap with the predicted cNLS regions (SFigure 9).

E3 ubiquitin-protein ligase RNF13 is a transmembrane protein localized to endosomes, with its targeting mediated by a dileucine motif (Leu311–Leu312) located in the cytoplasmic tail. Mutations in this motif have been associated with developmental and epileptic encephalopathies[12]. Although this motif matches the ELM regular expression (TRG_DiLeu_LyEn_5), it is not annotated in the database. SLiMMine successfully identifies this motif and additionally reports validated interactions with AP-3 complex sigma subunits[13], which are potential interacting partners for TRG_DiLeu_LyEn_5 class motifs.

**SNote 3: Regular expressions yielding overlapping motifs**

Sometimes the SLiMMine web server displays motifs slightly shifted, and thus fails to indicate them as ELM instances. For example, for the DOC_CYCLIN_RxL_1 and MOD_CK1_1 motifs of Cdc25A, the SLiMMine web server displays them as high-scoring hits, but not as known ELM instances, because the respective regular expressions yield multiple overlapping hits at the same region and only the first one was selected by the method each time (which does not necessarily match the instance defined in ELM).

Regular expressions are often defined such that a single motif can be detected multiple times within the same segment. Identification of overlapping motifs can occur either because the logical conditions in the regex allow overlapping matches, for instance, by flexibly defining sequence distances between two well-defined positions (for example, DOC_CYCLIN_RxL_1: SFigure 8, top), or because high-probability regular expressions, particularly in low-complexity regions, produce multiple overlapping motif hits (for example, MOD_CK1_1: SFigure 8, bottom). Several ELM classes (e.g., LIG_WD40_WDR5_VDV_2, MOD_GSK3_1, MOD_GlcNHglycan) exhibit this behavior, in several cases almost doubling the number of identified regex hits. While these "duplicated" regions rarely provide additional biological insight, they can inflate the total number of (potentially false) hits by hundreds of thousands. To prevent redundant listings of motifs shifted by only a few residues, we report only the first occurrence of each motif instance, and overlapping subsequent hits are not displayed.

**SNote 4: Protein-protein interaction data**

High-throughput experimental approaches, including affinity-purification mass spectrometry[13], complementary proximity-based labeling[14], yeast two-hybrid screening[15], and other large-scale techniques, have largely expanded the coverage of protein-protein interaction (PPI) data. Accordingly, databases storing PPI information have grown substantially over the past decades[16,17].

For the majority of these interactions, however, the molecular mechanism underlying the interaction remains unknown. Although BioGRID and IntAct collectively contain approximately 1,300,000 unique human PPIs annotated with physical association or higher evidence, only around 1,000 interactions have structural evidence from the PDB confirming that they are formed via domain–domain interactions (DDIs). Notably, BioGRID and IntAct do not incorporate PDB-derived structural data, which would increase the number of unique interactions by roughly 8,000[18].

Interestingly, approximately 2,000 experimentally validated unique human domain–motif interactions (DMIs) are available in ELM, LMPID, and PDB, which is twice the number of structurally resolved DDIs. While motif-based interactions are much more difficult to extrapolate across orthologs, DDIs can be extended more easily. By analyzing all DDIs from PDB (including all taxonomies) and mapping the corresponding PFAM domains to BioGRID and IntAct interactions, we identified ~20,000 PPIs that can reasonably be assumed to be formed via domain-domain interactions.

Thus, prior to SLiMMine, a total of ~23,000 PPIs (or ~31,000 when including PDB-derived interactions not stored in BIOGRID/IntAct) had some level of evidence about the molecular basis of the interaction. By defining potential interaction partners for all ELM classes and accurately predicting SLiMs across the human proteome, SLiMMine can inform on the molecular mechanisms and binding regions for an additional ~50,000 DMIs (SFigure 11).

**SNote 5: Dataset construction**

As with all supervised machine learning methods, SLiMMine is heavily influenced by its training set. ELM contains only a couple of thousand experimentally verified SLiM instances grouped into ~320 SLiM classes relevant to human proteins. While there are other sources of motifs, including annotated resources and high-throughput experiments, their reliability differs greatly. During training, we conducted experiments with larger datasets; however, including additional datasets likely introduced noise that prevented the neural networks from converging. Notably, even ELM motifs exhibit high heterogeneity: not all human motif instances are highly conserved, and some examples may not be fully disordered. On the one hand, this means that negative data gathered by collecting segments that seem unfavourable for forming a SLiM may contain positive examples, thereby hardening the training. On the other hand, even if protein embeddings capture a large number of features, the currently limited number of training examples leads neural networks to converge to somewhat different local optima, sometimes incorrect ones, leading to overfitting. For example, the training set included a large amount of negative data, in which protein localization prevents a motif from being functional (e.g., an extracellular integrin-binding motif cannot function on cytoplasmic proteins). Although we need such examples to increase the amount of training data, allowing the Neural Network to use features related to protein localization turned the predictor into a simple intracellular/extracellular predictor rather than learning other features. Even if we used cases with inappropriate localization for training, we could not include these examples in feature selection to prevent overfitting the NNs to localization information. We also had to separate intracellular and extracellular motifs, as these segments differ substantially. This is not surprising, as these proteins, subject to different environmental selection pressures, exhibit distinct characteristics; for example, disordered regions are much more prevalent within the cell. As the number of reliably verified SLiM examples increases, training similar methods may become easier. In some cases, the motif definition is somewhat permissive, leading to a high number of hits when scanning the proteome. In agreement, running SLiMMine on these less-defined instances may yield more false motifs (considering absolute values), even if specificity remains high.

**SNote 6: Pathogenic motifs**

Although SLiMMine was trained on human instances and is expected to work on mammalian motifs, we were interested in its predictive performance on proteins of more distant taxa.

Early E1A of Human adenovirus 5 (HAdV) is a widely studied effector protein (*E1A*; P03255). It is the first gene expressed after infection, since it can condition the cellular environment in favor of HAdV infection through concurrently interacting with multiple key host factors through SLiMs. Although E1A is known to mediate at least 32 primary interactions with host cell proteins, we here focused on how reliably SLiMMine can identify the twelve SLiM-mediated interactions depicted by King CR *et al.*[19] and/or stored in ELM[2]. Of the 5 ELM instances of E1A, (the C-terminal NLS, the C-terminal binding protein (CtBP)-binding PxDLS motif, two different Retinoblastoma-associated protein (Rb)-binding motifs and the BS69 (gene: *ZMYND11)* MYND domain-binding motif) SLiMMine could identify four (even though viral instances were not part of the training set). The motif binding into the deep groove formed between the A and B subunits of Rb was missed. SLiMMine could also identify a SUMO interacting motif (SIM; ELM class: LIG_SUMO_SIM_par_1) binding to the SUMO-conjugating enzyme UBC9, which fits the respective ELM regular expression but is not an ELM instance. Of the other six known motifs that do not fit any ELM regex, SLiMMine could identify the bipartite NLS binding to Qip1 (importin-α3) that has an unusually long linker between the two parts, the NES, an extended nuclear receptor box motif binding to the thyroid hormone receptor (TR), a motif binding to the Forkhead box protein K1 and K2 (FOXK1/K2) transcription factors and an amphipathic α-helix motif mimicking A-kinase anchoring proteins (AKAPs) that enables E1A to take control over protein kinase A (PKA). A motif binding to the TAZ2 domain of the CBP/p300 histone acetyl transferases (a candidate motif in the ELM resource) was missed by SLiMMine. In all, SLiMMine could successfully identify 10 of the 12 tested motifs of HAdV E1A, although it did not encounter viral proteins or motifs during training (SFigure 10). Notably, as class-independent identification yielded poor values on non-mammalian species, motifs without ELM class correspondence were assessed by using the start/end position of the peptide to calculate the SLiMMine score.

We further evaluated SLiMMine on all pathogenic proteins with at least one experimental motif instance in ELM. The positive set includes all (experimental) ELM instances in these proteins. The background set includes regex hits for all ELM classes within the same set of proteins that do not overlap with experimentally verified ones. The results show that SLiMMine predicted experimentally verified motifs with markedly higher score distribution than other regex hits. Notably, we expect multiple valid motifs among other regex hits (SFigure 14).

**SNote 7: Manual annotation of SLiMs**

During the curation of ELM classes we collected the following information (SData 1):

Accession: ELM instance identifier (format: ELMExxxxxx).

ELMIdentifier: ELM class name.

In (I)/Out (O): Broad localization in which the motif is functional: intracellular (inside; I) or extracellular (outside; O).

Binding domain ID: Broad definition/identifier of the domain that binds the motif.

All occurrences of the domain(s) in the proteome can bind the motif? (Y/N): Indicates whether all occurrences of the specified domain(s) in the human proteome are considered capable of binding the motif (Yes: Y; No: N).
Domain restricted to these proteins (regions if more): If not all occurrences of the specified domain(s) are considered to bind the motif, this column lists the UniProt accessions (ACs) of the specific human proteins containing the relevant binding domain(s).
Restriction confidence: The two previous columns define the motif-binding domains either by just domain type, or specific proteins when the domain type does not provide sufficient/unambiguous information. The confidence levels in this column signal if all relevant motif-binding proteins/domain occurrences have been precisely identified by the previous two columns. “Strict” means that to our knowledge all motif-binding domains are precisely covered and none of them are expected to be false positive hits. “Loosely defined” means that we listed all motif-binding domain occurrences that we know of and that are strongly suspected to bind the motif based on the respective ELM descriptions, but some of them do not have direct evidence for motif binding, just belong to the same family/subfamily of proteins that do have. In this case some true hits might be missed and false positives might occur. “Not defined” was added to the general motif definitions belonging to some of the largest domain families, such as SH3 and PDZ, where the binding specificity of many of the domains is not known and therefore a precise list of the domains binding to these general motif definitions is not possible to assemble. (Sometimes extra notes are provided).
Motif only found in proteins with this domain: Domain whose presence in the motif-containing protein is required for motif functionality (e.g., in cases of autoinhibition).
Motif only found in proteins with this GO process or activity: Gene Ontology (GO) terms describing biological processes or molecular activities associated with the functionality of the motif.
Motif restriction confidence: Confidence of the GO-based motif restriction based on annotations of proteins containing experimentally validated motif instances. Strict indicates that the GO term is consistently associated with motif-containing proteins, whereas Preferred indicates a less stringent association.
Domain-containing partner restricted to specific cellular location(s): Subcellular localization(s) to which the domain-containing counterpart of the motif-mediated interaction can be restricted.
Motifs always in TM proteins? (Y/N): Indicates whether the motif is restricted to transmembrane proteins (Yes: Y; No: N).
Localization of partners with motifs (specific): Specific Gene Ontology terms describing the subcellular localization in which the motif is functional. Because these annotations are relatively specific, the corresponding GO term may not be associated with every experimentally validated motif instance.
Localization of partners with motifs (less specific): Broader Gene Ontology terms describing the subcellular localization in which the motif is functional.

**SNote 8: Motif classes without any positive experimental instance**

Of the 319 ELM motif classes manually refined for SLiMMine, 290 contained at least one suitable positive instance and could therefore be included in benchmarking and predictor construction. For the remaining 29 classes, no suitable positive instance could be included in the human benchmark set. These classes comprise motifs with no instances stored in ELM, instances that could not be mapped to the current human reference proteome, instances available exclusively from distantly related mammalian species or pathogens, and overlapping motifs that could not be evaluated independently. These classes may nevertheless represent functional motifs in humans and were therefore retained in the refined SLiMMine annotations. The affected classes and the reasons preventing their inclusion in benchmarking are listed below.

- **No instance stored in ELM:** DEG_Cend_FEM1AC_1, DEG_Nend_Nbox_1, DEG_Nend_UBRbox_1, DEG_Nend_UBRbox_2, DEG_Nend_UBRbox_3, LIG_BIR_II_1, MOD_Cter_Amidation, MOD_NEK2_2.
- **Instances could not be mapped to the current human reference proteome:** DEG_Cend_KLHDC2_1, LIG_CSK_EPIYA_1, LIG_BIR_III_1, LIG_Integrin_KxxGD_FGGC_5, LIG_Integrin_RGDW_4, LIG_Mtr4_Trf4_2, LIG_MYND_1, LIG_SH2_STAT6.
- **Available instances are exclusively from distantly related mammalian species:** LIG_WRPW_2, MOD_PK_1, TRG_Cilium_Arf4_1.
- **Only pathogen/hijacking instances are available:** DOC_USP7_MATH_2, DOC_MAPK_GRA24_9, LIG_LYPXL_L_2, LIG_LYPXL_SIV_4, TRG_PTS2, DOC_RSK_DDVF_1.
- **Overlapping motifs, as discussed in SNote 2:** DOC_CYCLIN_RevRxL_6, DOC_MIT_MIM_1, LIG_LIR_LC3C_4, LIG_RBL1_LxSxE_2.

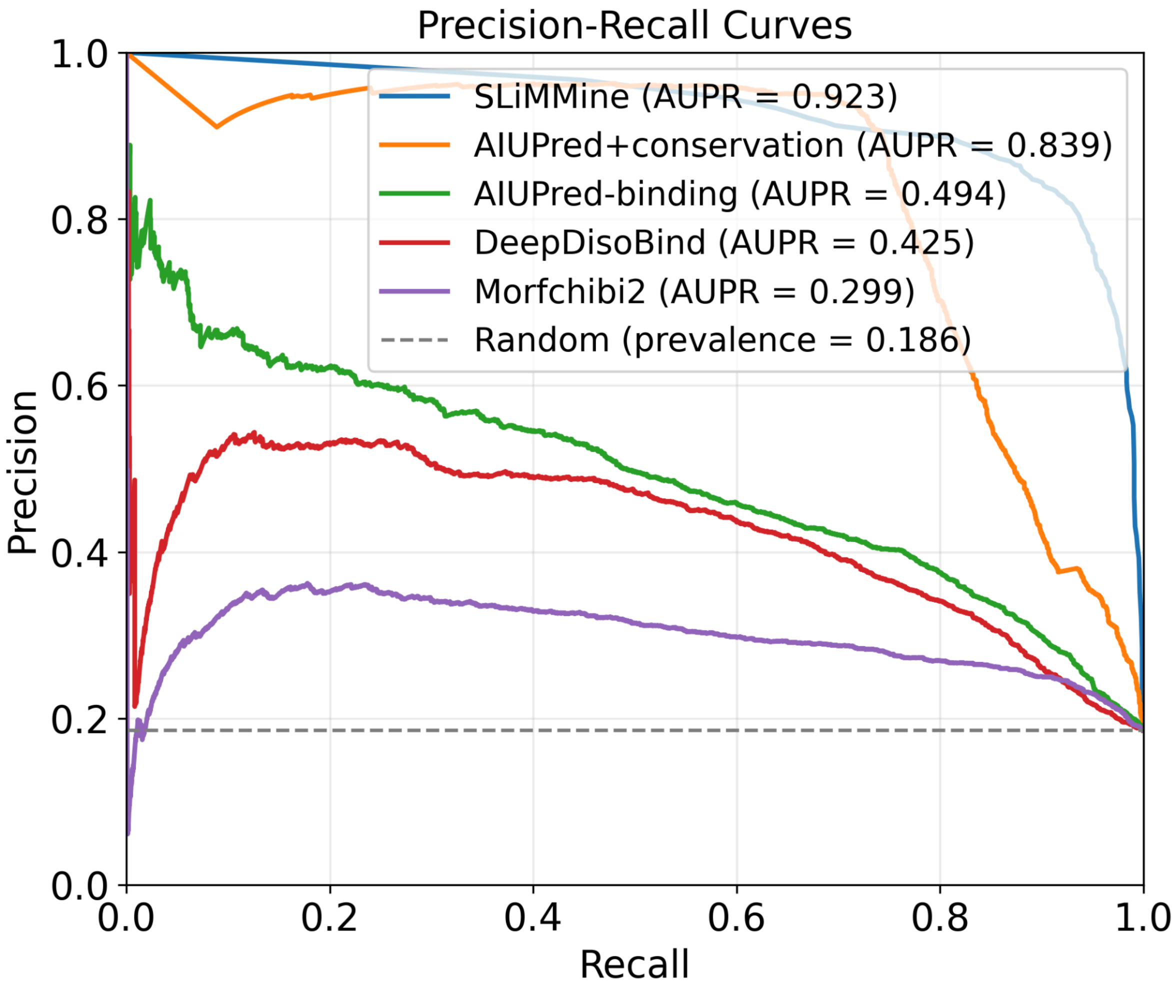


***SFigure 1: Comparison of precision-recall curve of SLiMMine and other methods.*** *Precision–recall curves and Area Under the Precision–Recall Curve (AUPR) values are shown for SLiMMine, AIUPred-binding, DeepDisoBind, MorfChibi2, and the traditional conservation- and disorder-based prediction approach.*

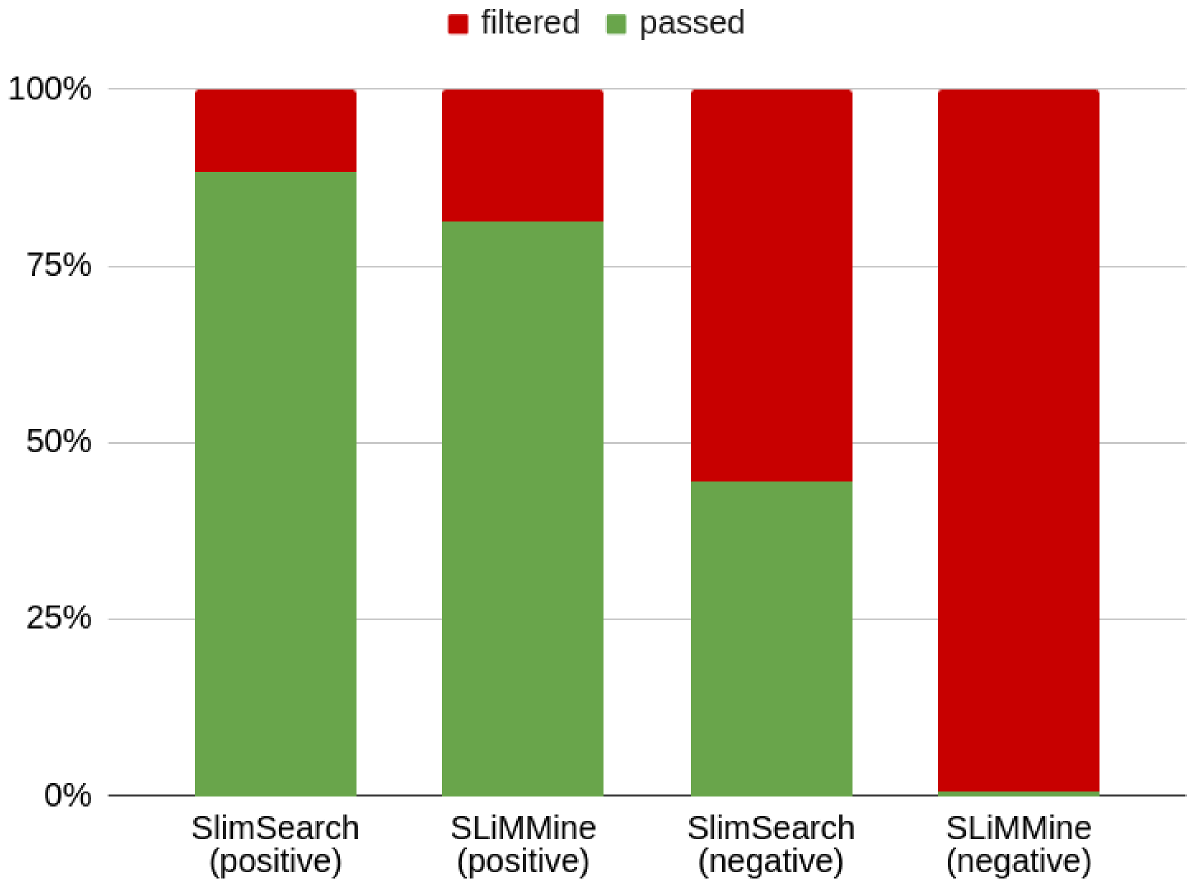


***SFigure 2: Comparison of SLiMSearch filters and SLiMMine predictions.*** *The proportions of motifs from the positive dataset and segments from the negative dataset passing (green) or failing (red) the SLiMSearch contextual filters or the SLiMMine prediction and contextual filters are shown. For SLiMMine, predictions with a probability >0.5 without any (problematic) filtering were considered positive. Cases for which no SLiMSearch result was available are excluded from the comparison.*

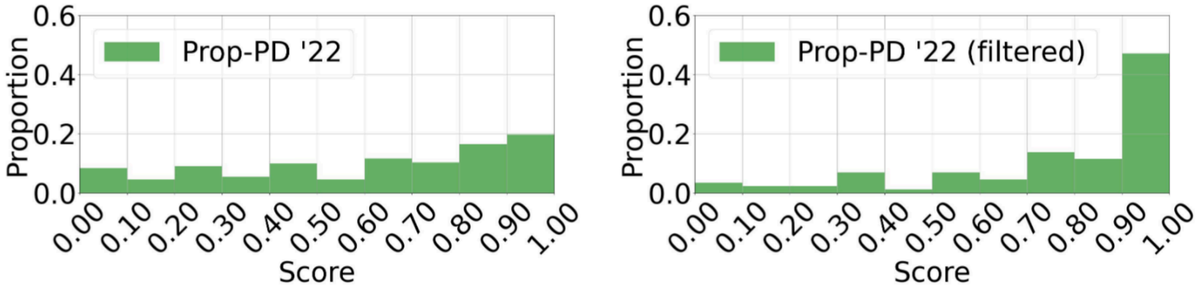


***SFigure 3: SLiMMine performance on Prop-PD dataset.*** *A) Distribution of SLiMMine scores on high-confidence Prop-PD peptide hits that do not overlap with ELM instances B) Distribution of SLiMMine scores when only conserved high-confidence Prop-PD hits are retained.*

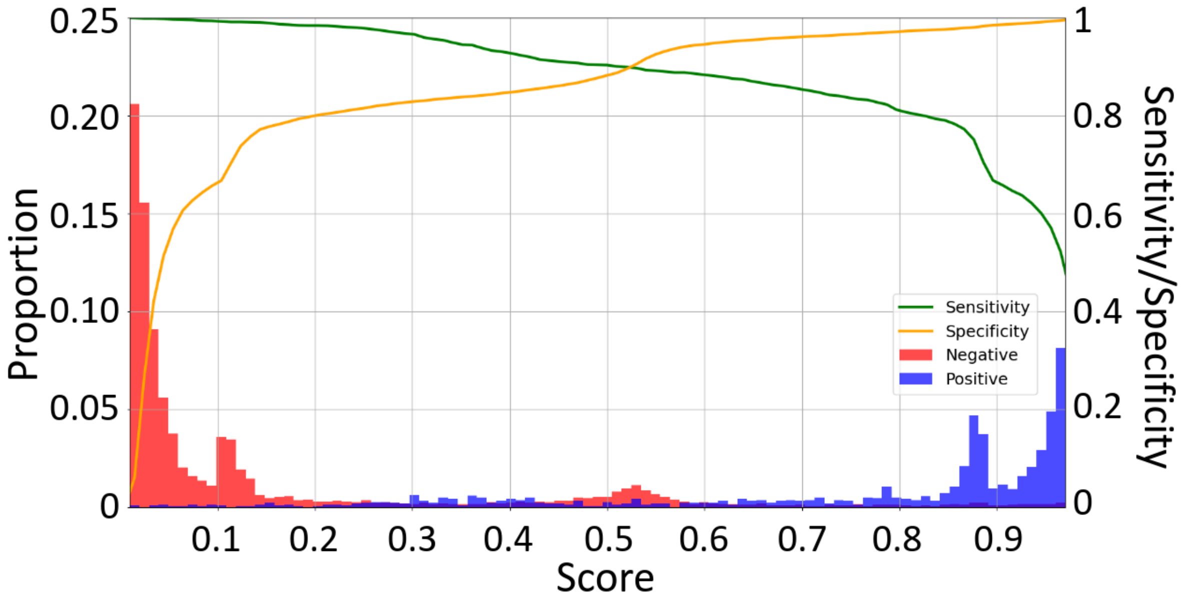


***SFigure 4: The performance parameters of SLiMMine on the benchmark set.*** *Distribution of prediction scores on the benchmark set (bars). Lines indicate the sensitivity and specificity at varying score thresholds.*

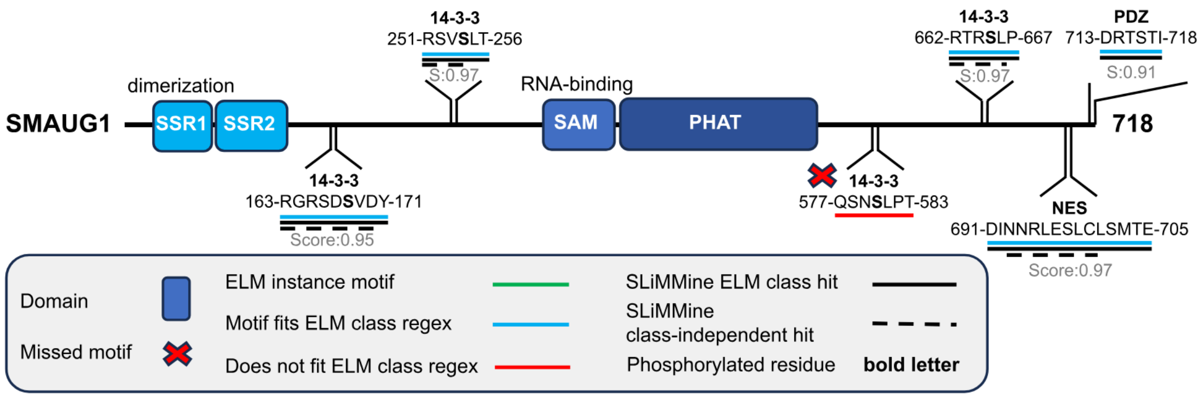


***SFigure 5: SLiMMine finds almost all experimentally validated motifs of Smaug1.*** *The known domains (boxes) and motifs (sequence bits) are indicated on the domain map of the Smaug1 RNA-binding protein. For each of the six validated motifs, their binding domains/motif names, sequences, residue boundaries, ELM status (underscoring in different colors) and SLiMMine predictions (underscoring with a simple (ELM class predictions) or dashed (class-independent predictions) black line) are indicated along with the respective SLiMMine scores. The phosphorylated S/T residues of the 14-3-3 binding motifs are highlighted in bold.*

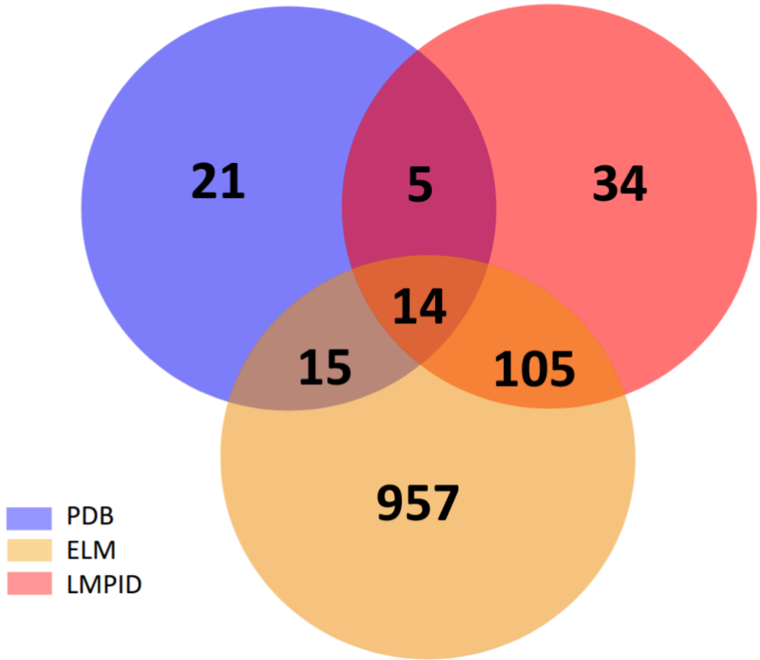


***SFigure 6: SLiMMine class-independent motif prediction finds multiple experimentally verified motifs.*** *Venn diagram showing the overlaps between the class-independent hits of SLiMMine and motifs found in PDB, LMPID and ELM.*

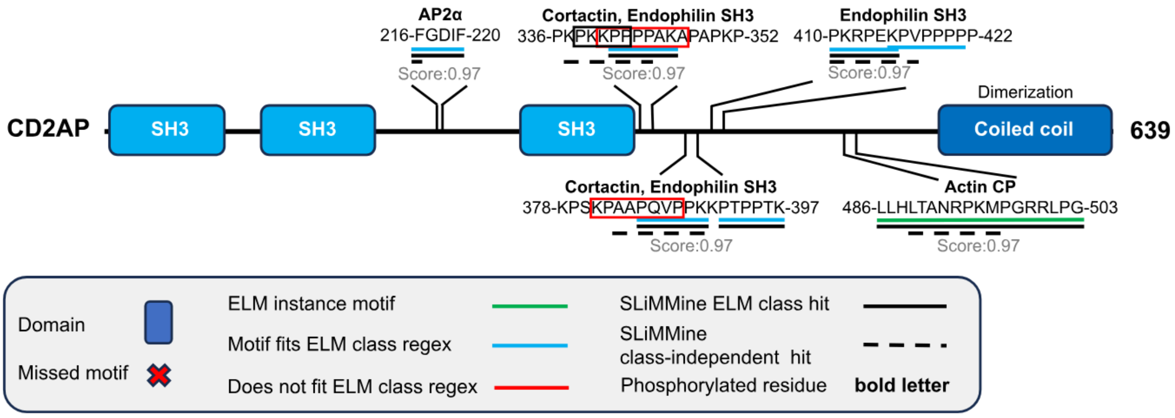


***SFigure 7: SLiMMine successfully identifies most known motifs of the human CD2AP.*** *The domains (blue boxes) and SLiMs (sequence bits) are indicated on the domain maps of CD2AP. For each known motif, the respective binding domains/motif names, sequences, residue boundaries, ELM status (underscoring in different colors) and SLiMMine predictions (underscoring with black simple (ELM class predictions) and/or dashed (class-independent predictions) line) are indicated along with the respective SLiMMine Score (S). In CD2AP, three Pro-rich peptides were shown to bind to the Cortactin and Endophilin-A1 SH3 domains, but the exact binding sites were not determined. ELM-annotated SH3-binding motifs (classes: LIG_SH3_1/2) are highlighted with blue underscoring in these peptides, the likely Endophilin-A1 SH3-binding site is highlighted with a black box, and the likely Cortactin SH3-binding sites with red boxes.*

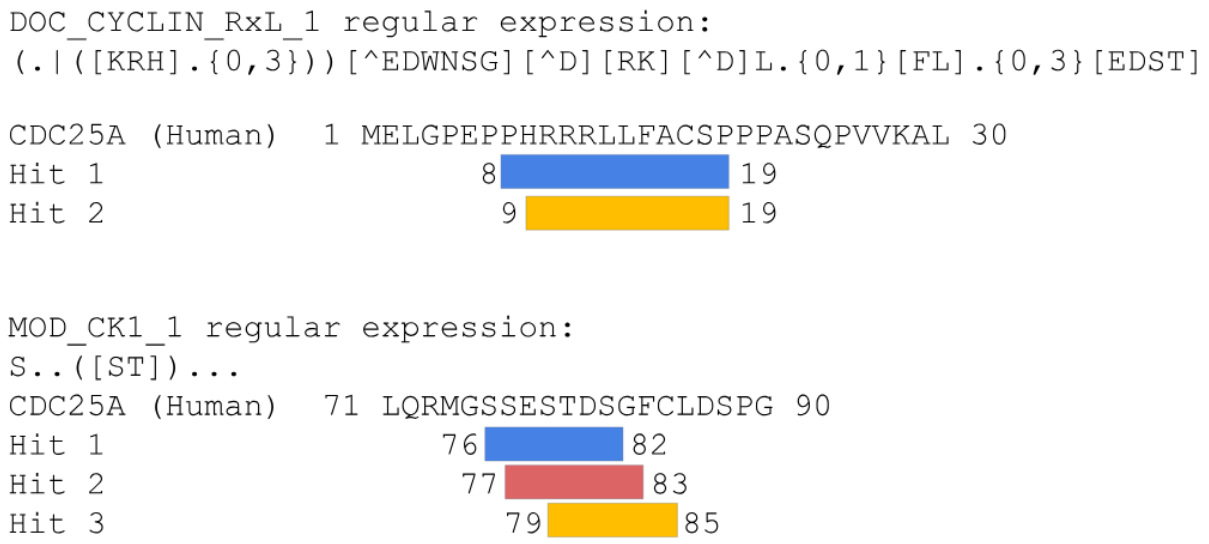


***SFigure 8: Overlapping and shifted motif detection may affect the visualization of ELM instances in SLiMMine.*** *The regular expressions of the Cyclin-docking motif and CK1 phosphorylation site from ELM yield overlapping motifs in Cdc25A.*

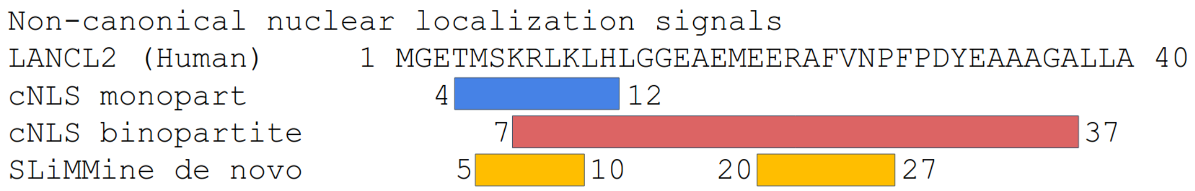


***SFigure 9: SLiMMine finds the nuclear localization signal in LanC-like protein 2 as a class-independent motif.*** *The nuclear localization signal in LanC-like protein 2 (LANCL2) does not fit the ELM definition. Although the exact motif start and end positions have not been experimentally verified, class-independent motifs from SLiMMine overlap with those predicted by cNLS.*

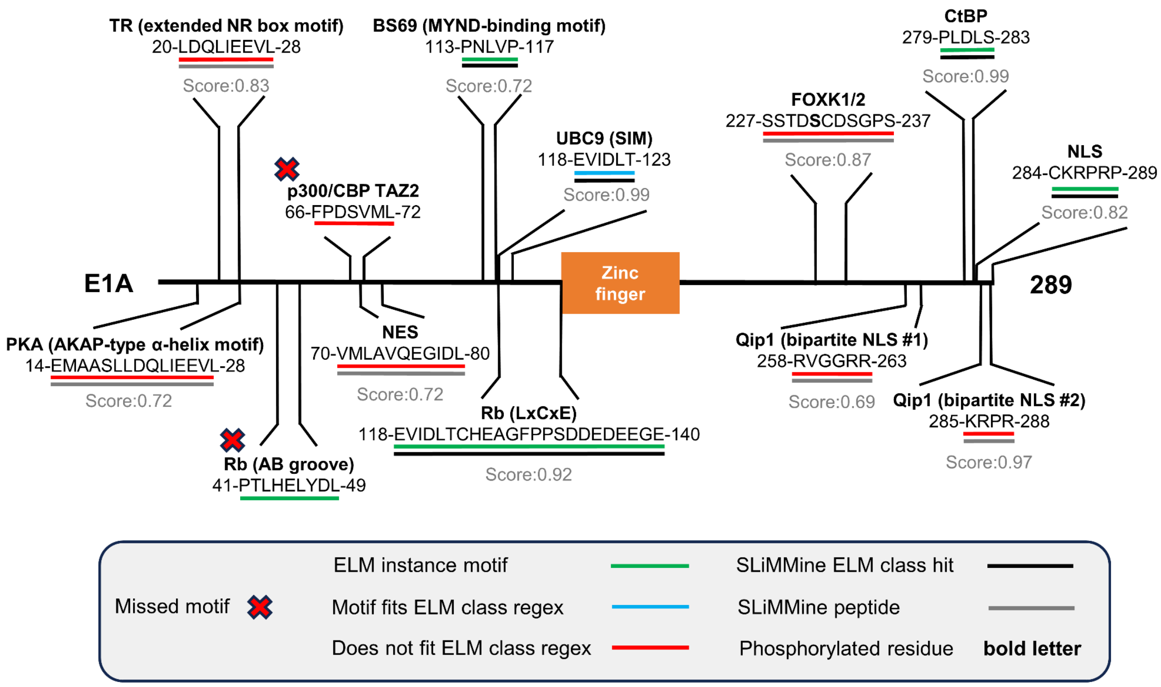


***SFigure 10: SLiMMine successfully identifies the motifs employed by the viral Early E1A protein.*** *Experimentally validated motifs (sequence bits) are indicated on the domain map of the E1A protein of HAdV. For each validated motif, the binding partner/motif name, sequences, residue boundaries, ELM status (underscoring in different colors) and SLiMMine predictions (underscoring with black (ELM class predictions) or grey (prediction based on the start/end position of the peptide)) are indicated along with the respective SLiMMine scores.*

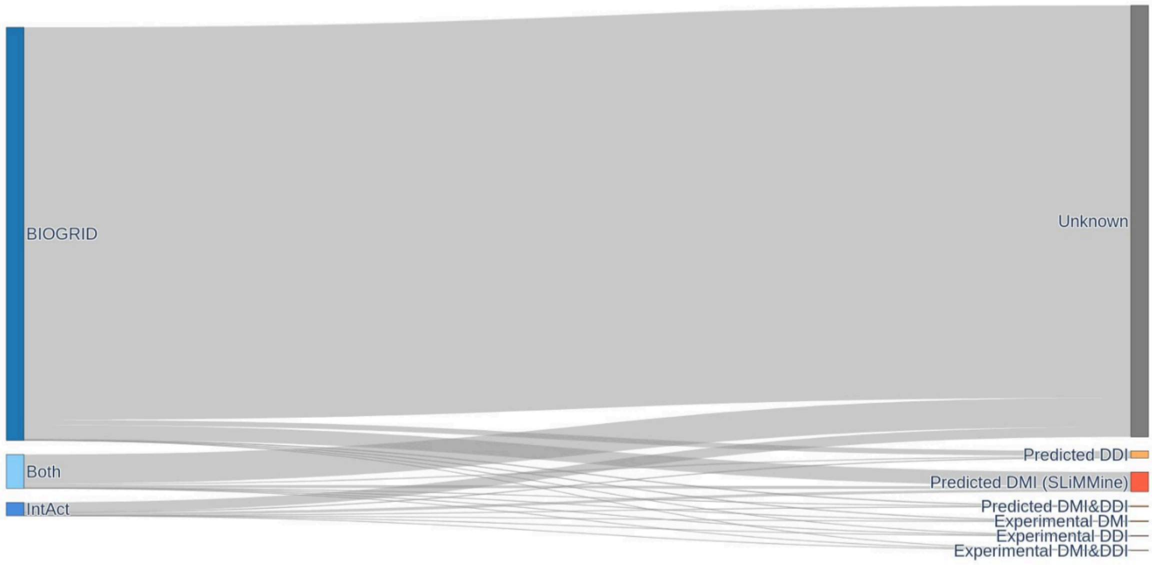


***SFigure 11: SLiMMine predictions explain a non-negligible fraction of validated human PPIs.*** *Experimentally verified PPIs from BIOGRID and IntAct fall into different interaction types regarding the interacting protein modules. Experimental Domain Motif Interactions (DMIs) are retrieved from ELM, LMPID and 3did (PDB), experimental Domain Domain Interactions are retrieved from 3did (PDB), predicted DDIs are interactions between proteins with PFAM identifiers that were detected to interact in PDB structures (via 3did), predicted DMIs come from linking SLiMMine predictions to the PPIs.*

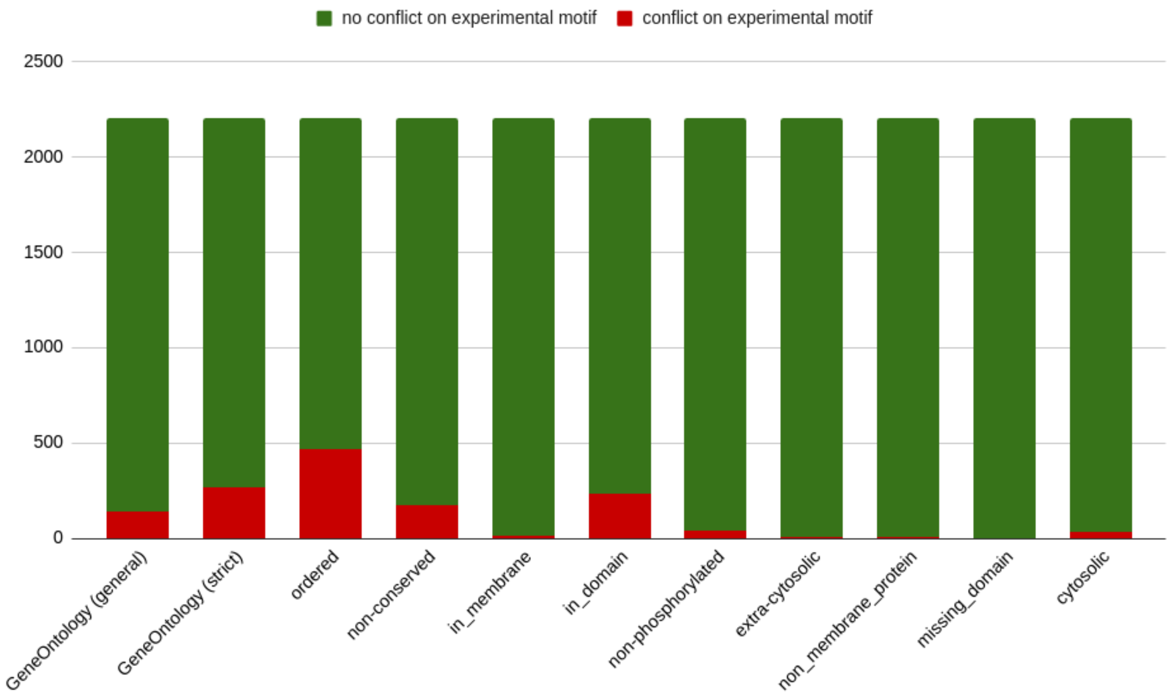


***SFigure 12: The effect of contextual contradictions on experimentally verified motifs.*** *The proportion of falsely excluded experimentally verified SLiMs (red) in the training set. The SLiMMine web server uses the same filters.*

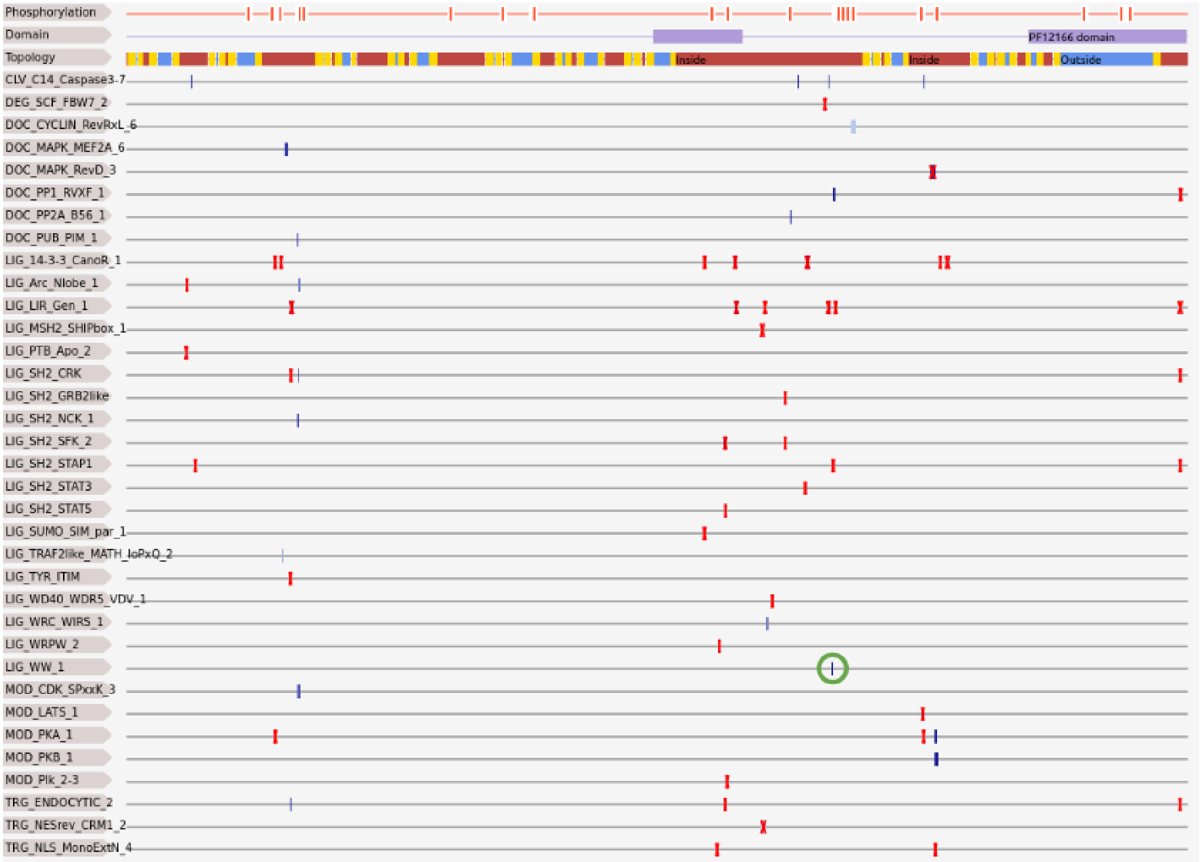


***SFigure 13: Context filters work in the Piezo2 receptor.*** *The snapshot of the SLiMMine webserver shows the domains, topology and the predicted well-defined CLV, DEG, DOC and LIG type motifs of the giant Piezo2 transmembrane receptor (UniProt AC: Q9H5I5). While most of the predicted motifs are filtered out by our pre-defined contextual filters (motifs with red crosses), the WW domain-binding motif detected to bind to YAP1 and NEDD4 WW domains in a ProP-PD screen remained as a high-scoring hit (1828-PPSY-1831; score: 0.97; highlighted in a green circle).*

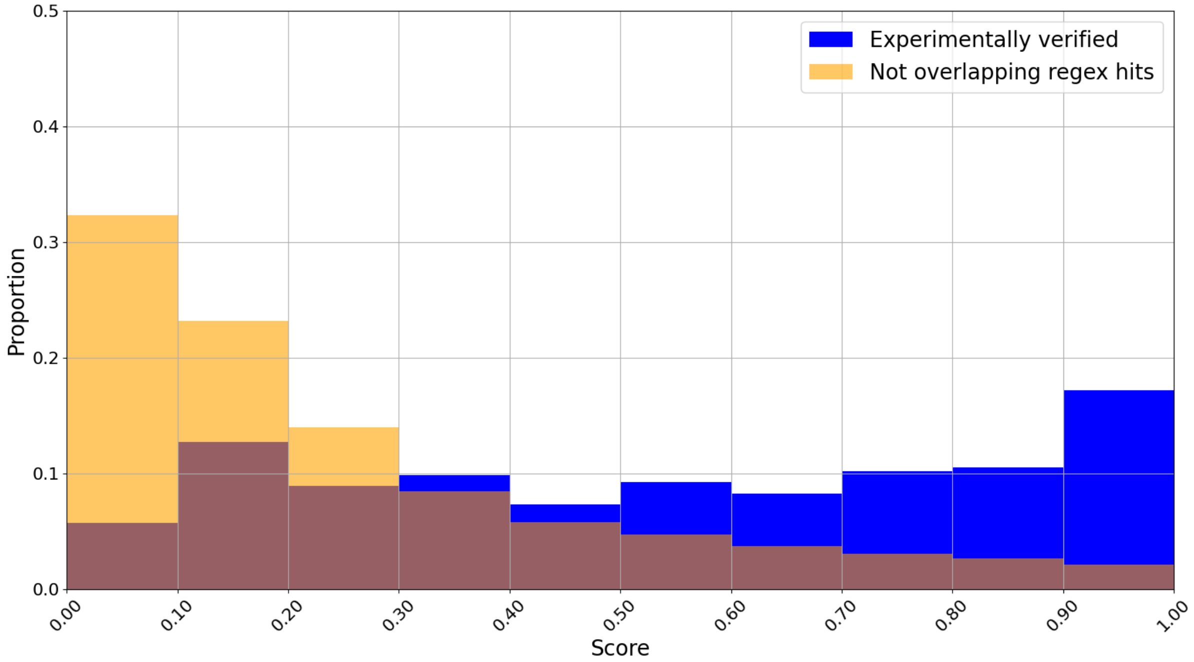


***SFigure 14 SLiMMine performance on motifs of pathogens.*** *SLiMMine scores for pathogenic ELM instances (blue) (only human instances were used for method training) and all other regular expression hits detected for the same protein pool that were not overlapping with the experimental ones (yellow).*

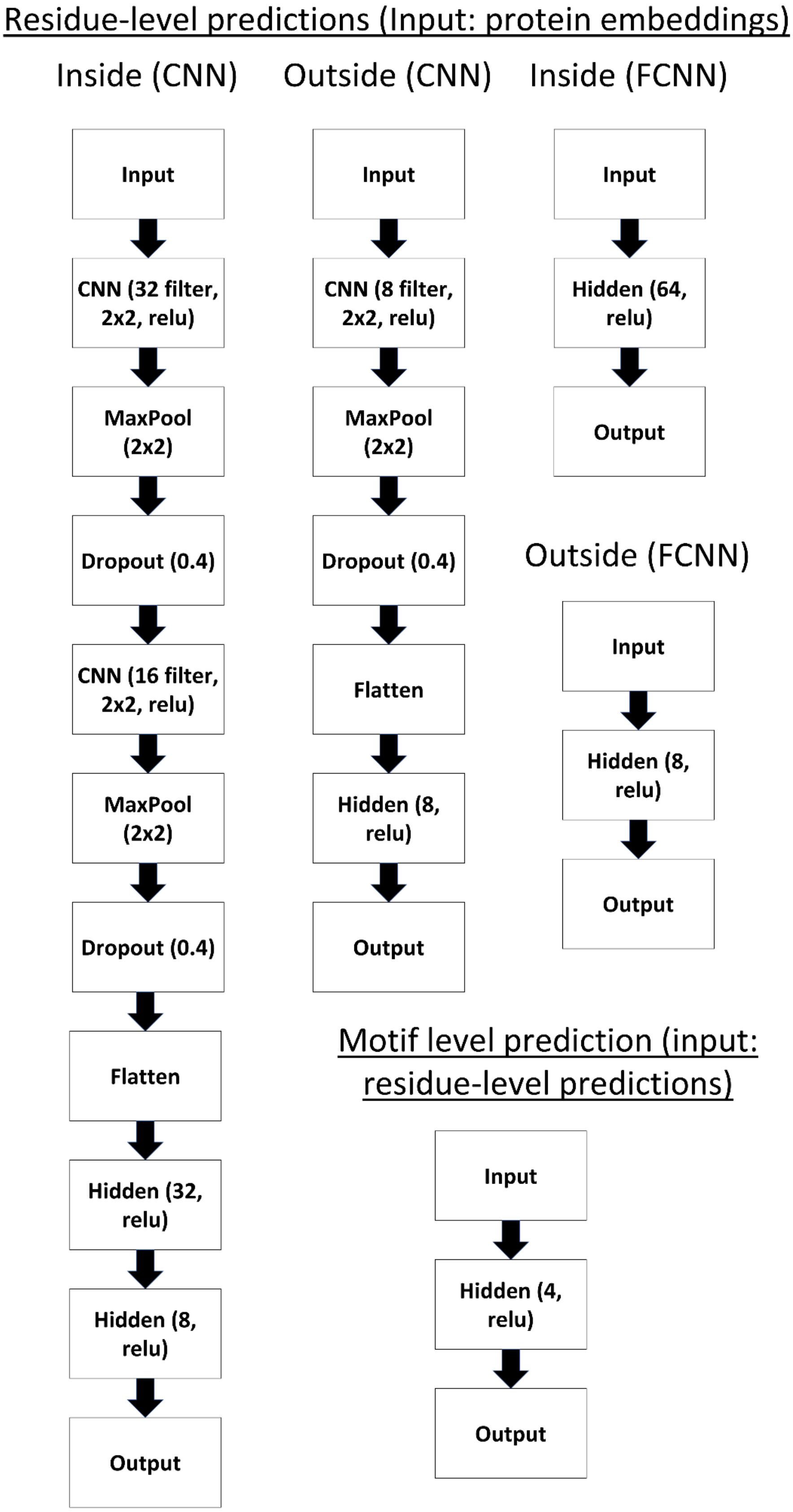


***SFigure 15: The Neural Network architectures of SLiMMine.***

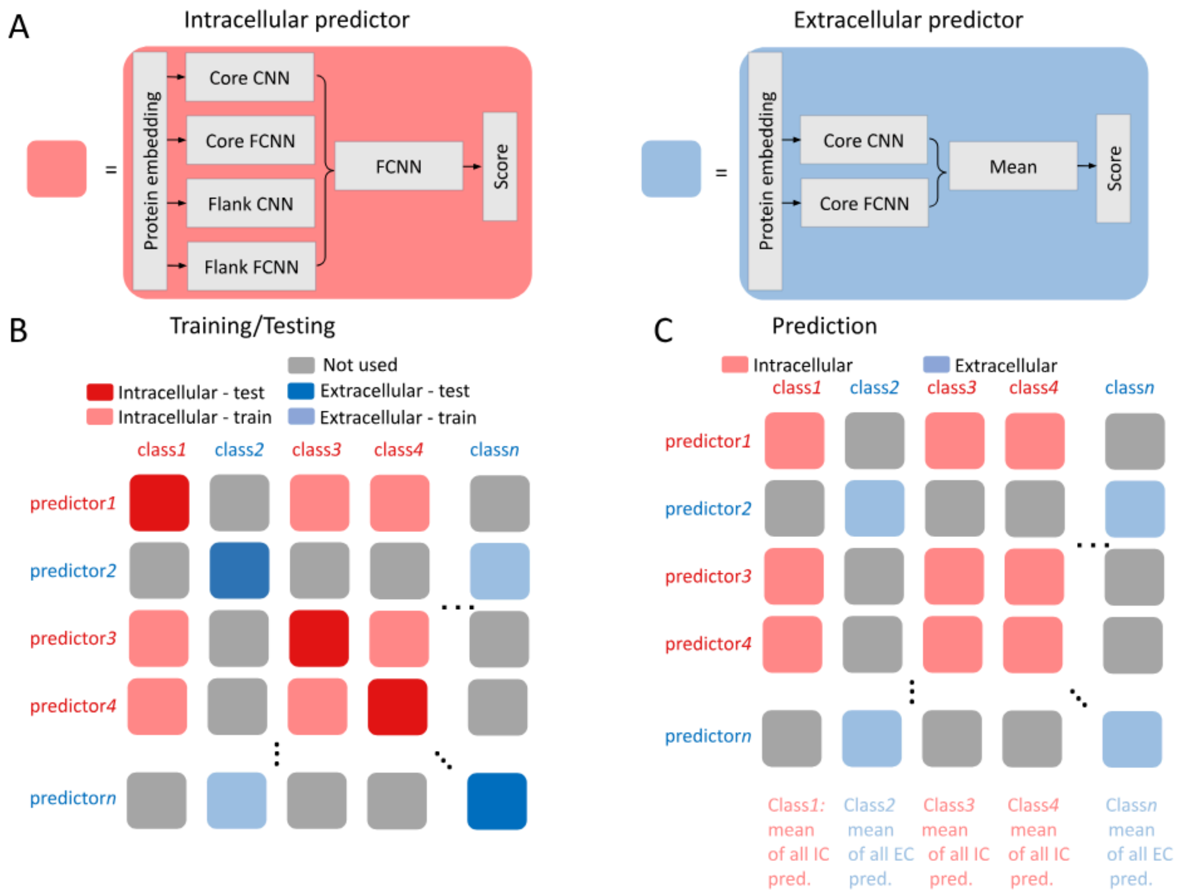


***SFigure 16: Use of predictors during benchmarking and proteome-wide prediction.*** *A) Schematic representation of the intracellular and extracellular predictors. B) Selection of predictors during the training and benchmarking of SLiMMine. C) Selection of predictors for predicting motifs across the human proteome.*
